# Characterising anthelmintic resistance against benzimidazoles and macrocyclic lactones in gastrointestinal nematode populations of dairy cattle

**DOI:** 10.1101/2025.09.14.676042

**Authors:** Paul Campbell, Jennifer Mcintyre, Kerry O’Neill, Andrew Forbes, Kathryn Ellis, Roz Laing

**Affiliations:** School of Biodiversity, One Health and Veterinary Medicine, University of Glasgow, Glasgow, UK

**Keywords:** Gastrointestinal nematodes, Egg hatch test, Anthelmintic resistance, Benzimidazole, Surveillance

## Abstract

Anthelmintic resistance in gastrointestinal nematode populations is endemic across all grazing livestock production systems, particularly in small ruminants, but also in cattle. In this study resistance in *Cooperia oncophora* and *Ostertagia ostertagi* to macrocyclic lactones and benzimidazoles (BZ) in Scotland was assessed. The faecal egg count reduction test (FECRT) remains the primary tool for assessing anthelmintic resistance in the field; however, differing methodologies and recent guideline updates complicate the interpretation of results across studies. Statistical approaches, such as Bayesian methods used by *eggCounts* and *bayescount*, produce varying confidence intervals, which further influence conclusions.

The ability of the egg hatch test to detect anthelmintic resistance was evaluated under field conditions, and the EC_50_ values obtained enabled assessment of the BZ efficacy at both the farm and GIN species levels, without the need for treatment. Mixed amplicon sequencing was applied, and nemabiome analysis confirmed that *C. oncophora* and *Os. ostertagi* were the predominant species. Sequencing of the *beta-tubulin isotype-1* gene in strongyle communities identified resistance alleles in multiple species.

The high EC values and elevated frequencies of BZ-resistance alleles on some farms emphasise the importance of *in vitro* assays and molecular diagnostics for farm-level resistance management. The detection of resistance against both benzimidazoles and macrocyclic lactones on certain farms highlights the urgency of implementing sustainable control strategies. Although the data presented here are from Scotland, given the high rate of animal movement and similar patterns of anthelmintic usage in the UK, these findings are likely to be relevant across much of the UK.

**Highlights:**

- Resistance to MLs and BZs was detected in GIN populations of cattle
- Multidrug resistance found in Scottish GIN populations of cattle
- Egg hatch test results suggest, and sequencing confirms high BZ-resistance allele frequencies

## 1. Introduction

Gastrointestinal nematode (GIN) infections are a major cause of reduced productivity and adverse effects on animal health and welfare. Broad-spectrum anthelmintic drugs have been the backbone of parasite management in livestock for 60 years; however, their continuous use has driven the selection of drug-resistant populations worldwide. The high prevalence and rapid emergence of anthelmintic resistance, including multidrug resistance, in the sheep industry have been the focus of extensive research.

In contrast, the ostensibly lower rate of anthelmintic resistance detection in cattle in temperate regions may partly be due to diagnostic challenges, as faecal egg counts are generally low. Egg counts are considerably less reflective of the actual worm burden compared with sheep, making detection of resistance using the faecal egg count reduction test (FECRT) inherently more uncertain. Simultaneous resistance to all three drug classes, benzimidazoles (BZ), macrocyclic lactones (ML), and imidazothiazoles, has now been identified in the two main GIN species infecting cattle: *Cooperia oncophora* and *Ostertagia ostertagi* in New Zealand (Sauermann et al., 2024).

Reports of anthelmintic resistance in UK cattle remain limited and do not currently suggest a severe problem. Nevertheless, UK studies (Bartley et al., 2012; Geurden et al., 2015; McArthur et al., 2011; Stafford and Coles, 1999) have indicated low to moderate levels of ML resistance. By contrast, resistance to BZs appears rare, with only one resistant population identified to date (Bartley et al., 2021). However, the number of studies conducted and farms surveyed is small, and current data are insufficient to provide a representative national picture (Rose Vineer et al., 2020).

Grazing cattle are particularly susceptible to GIN infection during their first two grazing seasons, after which they typically develop immunity and maintain? low FECs. Youngstock, however, shed relatively more eggs and are therefore the most suitable age group for evaluating anthelmintic efficacy using the FECRT. In organic systems, ML use is restricted, and regulations strongly discourage whole-group anthelmintic treatments, instead promoting evidence-based treatment approaches such as targeted-selective-treatment strategies. Organic producers primarily use BZ or levamisole (LEV) products to treat GIN infections, therefore nematode populations on these farms should have little or no exposure to MLs.

To address the knowledge gap regarding anthelmintic resistance in Scottish cattle, we conducted a study investigating the prevalence of resistance in first-grazing-season dairy calves. The FECRT was used to evaluate the efficacy of the most commonly used anthelmintics: ivermectin, moxidectin, and fenbendazole, with the egg hatch test (EHT) also used as an alternative measure of BZ resistance. In addition, mixed amplicon sequencing was employed to characterise GIN species composition and to determine the frequency of genetic markers associated with BZ and LEV resistance.

## 2. Methods

### 2.1. Ethics statement

The University of Glasgow MVLS College Ethics Committee (Project No: 200210097) approved all research procedures involving animal use.

### 2.2. Selection of farms

All 14 farms were selected on the following criteria: located in Scotland; herd size ≥30 first-grazing-season (FGS) calves, no anthelmintic treatment administered during the current grazing season; and a minimum of two months of grazing before sample collection. Additionally, all farms were required to complete a livestock management survey. The four farms participating in the FECRT were also required to have a cattle crush and handling system available.

All farms were offered free FEC analyses and evaluation of anthelmintic efficacy by FECRT. All farms participating in the FECRT were conventionally managed whereas those participating only in the EHT were all organically managed

### 2.3. Farm survey

A questionnaire was completed during a semi-structured interview, collecting demographic data, information on pasture management and anthelmintic usage, and experiences with faecal egg counts and other helminth infections. For more details, see Chapter 2, Section 2.5 and Appendix X.

### 2.4. Faecal egg count reduction test protocol

All animals were turned out to pasture in May 2023. From eight weeks post-turnout, faecal egg counts (FECs), were monitored fortnightly until the group mean FEC reached ∼100 eggs per gram (EPG). In addition, bovine lungworm (*Dictyocaulus viviparus*) larvae in faeces and body condition were measured regularly. On the day of treatment (Day 0) on each farm, 15 calves were randomly allocated to a treatment group and received either fenbendazole (Panacur® 10 % Oral Suspension; MSD Animal Health) *per os*, subcutaneous ivermectin (IVOMEC® Classic Injection for Cattle and Sheep; Boehringer Ingelheim), or moxidectin (Cydectin® 10 % LA Solution for Injection; Zoetis) by subcutaneous ear injection, at the manufacturers’ recommended dose rates of 7.5, 0.2, and 1.0 mg/kg of body weight respectively. Calves were either individually weighed or their weight estimated using a dairy calf weight band (AHDB), with weights ranging from 187 to 239 kg. Dose calculations were performed by the researchers, and anthelmintics were administered accordingly, with each dosage rounded to the nearest practical measure based on the formulation: 1 ml for fenbendazole, 0.1 ml for ivermectin (IVM), and 0.05 ml for moxidectin (MOX).

Faecal samples were collected *per rectum* from all animals on day 0 (pre-treatment) and day 14 (post-treatment). Samples were sealed immediately after collection and transported to the University of Glasgow on the same day. Aliquots of faecal material were stored at 4 ^°^C for FEC analysis, while samples for egg isolation and coproculture were stored at room temperature and processed within 24 hrs of collection. On all farms, calves from the different treatment groups were grazed together on the same pastures until day 14.

### 2.5. Faecal egg count

A faecal egg count was performed on every individual animal sample using a modified salt flotation technique as described in Jackson, 1974, with a detection limit and multiplication factor of 1 egg per gram (epg). Briefly, faecal samples (3g) were homogenised in 10 ml of water per gram of faeces suspension and passed through a 1 mm sieve, followed by rinsing with 5 ml of water to remove debris. The filtrate was then transferred to polyallomer centrifuge tubes and centrifuged at 200 g for 5 minutes. The supernatant was discarded, and the pellet resuspended in saturated salt solution (SSS) (specific gravity of 1.2), vortexed, and centrifuged at 200 g for 10 minutes. The meniscus was then isolated using haemostat clips, and the suspension transferred to a cuvette, which was then completely filled with SSS. Cuvettes were then read after waiting a minimum of five minutes, and GIN eggs were identified morphologically and counted as either strongyle-type or *Nematodirus* spp.

For each individual animal, the number of strongyle eggs present pre- and post-treatment was used to calculate the percentage reduction in FEC, thereby estimating anthelmintic efficacy. In accordance with the revised FECRT guidelines (Kaplan et al., 2023), a minimum mean of 40 strongyle-type eggs per animal pre-treatment was required for reliable assessment. This threshold was achieved by performing one FEC per individual; however, to facilitate species-specific faecal egg count reduction calculations, it was estimated that a total of three FECs were needed per sample.

### 2.6. Egg hatch Test

Egg hatch tests on farms participating in the FECRT were conducted using eggs pooled from all pre-treatment animals. For the organic farms, eggs were pooled from faeces collected from pasture using the methodology described in Chapter 2, section 2.2.

Eggs were isolated from pooled fresh faeces by sieving, centrifugation and flotation in SSS. Briefly, 300g of faeces were mixed with tap water, passed through 500 µm and 210 µm sieves, and centrifuged at 2,500 rpm for 5 minutes. The supernatant was discarded, and kaolin was added to the pellet, and the mixture was vortexed before being resuspended in the SSS.

After centrifugation at 1,000 rpm for 10 minutes, the polyallomer tubes were clamped to isolate the eggs, which were then collected in a 38 µm sieve, rinsed thoroughly with deionised water, and examined microscopically to confirm that embryonation had not yet begun.

Each sample was tested in triplicate using six concentrations of thiabendazole (TBZ) (0.01, 0.025, 0.05, 0.1, 0.25 and 0.5 μg TBZ / ml) and a negative control (no drug, 0.5 % DMSO), also in triplicate. The EHT was performed following the protocol described by von Samson-Himmelstjerna et al., 2009. After 48 hours, the test was terminated with Lugol’s iodine stain, the contents of each well were transferred to a Petri dish, and all eggs and larvae were counted. The contents from each well were then pooled by TBZ concentration for species identification. Up to 94 eggs/larvae were identified to species level for each TBZ concentration as described in Chapter 4, Section 2.5.

### 2.7. Coproculture and species identification by PCR

Pooled larval coprocultures from each pre- and post-treatment group were prepared by hand-mixing approximately 300g of faeces with vermiculite to form a well-aerated, uniform paste-like consistency. The coprocultures were incubated at 27 ^°^C for 14 days and sprayed with water as needed to maintain adequate moisture for L3 development. After incubation, larvae were harvested using a modified Baermann technique as described in Roberts and O’Sullivan, 1950; pooled aliquots of L_3_ in ddH_2_O were stored at –80 ^°^C.

Crude lysates were prepared from single strongyle eggs or larvae in 96-well plates using a modified proteinase K lysis reaction for individual strongyles identified by PCR. Briefly, a 100x stock solution of lysis buffer was made as follows: 1,000 µl DirectPCR Lysis Reagent (Cell) (Viagen Biotech), 50 µl 1M DTT (Invitrogen), and 10 µl Proteinase K (100 mg/ml) (Invitrogen). A 10 µl was aliquot of this buffer was dispensed per well of a 96-well PCR plate. Using a stereomicroscope, individual strongyle eggs or larvae were transferred into each well in a volume of ≤1 µl. Lysates were incubated at 60 ^°^C for 2 h, followed by 85 ^°^C for 45 minutes to denature the Proteinase K. Crude lysates were diluted 1:20 with nuclease-free water.

For each pooled pre-/post-sample, a minimum of 94 recovered larvae/eggs were identified to species level by PCR targeting the ITS2 region. A multiplex PCR method (Bisset et al., 2014) was employed, using primers designed to amplify the strongyle ITS2 region and species-specific regions for GIN species of interest (see Appendix File 2; Table S2). The reaction set included primers targeting *Haemonchus contortus*, *Os. ostertagi*, *C. oncophora*, *Oesophagostomum venulosum*, and *Trichostrongylus. axei*. Multiplex PCRs were performed in 96-well plates with 94 individual worm lysates, one DNA-negative control (no genomic DNA template), and one lysis-negative control (lysate without larva).

### 2.8. Genomic DNA isolation and amplicon sequencing library preparation

Genomic DNA from pools of 3,000 L_3_ was isolated using the Monarch® Spin gDNA Extraction Kit (T3010S) following the manufacturer’s instructions, with a final elution in 30 μl of buffer (10 mm Tris-HCl, pH = 9.0, 0.1 mm EDTA). The isolated gDNA was normalised to a concentration of 25 ng/μl, and all samples were stored at 4 °C.

Anthelmintic resistance mixed amplicon sequencing libraries were generated by individual PCR amplification of the loci as described in Chapter 3, Section 2.4. The primers for the ITS2 and *beta-tubulin isotype-1* loci were modified to include pan-nematode primer pairs developed by Avramenko et al., 2019, 2015. For the complete list of primers and thermocycling parameters used, see Appendix X. The PCR reactions (50 μl total volume) contained: 10 μl 5x Phusion GC buffer (New England Biolabs), 1 μl of 10 mM dNTPs, 2.5 μl of 10 μM of each forward and reverse primer (or primer mix), 0.5 µl of Phusion DNA Polymerase, 1 μl of gDNA, and 32.5 μl of nuclease-free water. All PCR steps were performed following best practices to minimise aerosol formation, including the use of filter pipette tips, working in a PCR cabinet, and sealing PCR plates with adhesive seals.

The unindexed amplicon libraries were submitted for Illumina MiSeq amplicon sequencing. Stage-2 indexing PCRs, library quantification, normalisation, pooling, and denaturing were performed according to Illumina’s 16S Metagenomic Sequencing Library Preparation (Illumina Inc., USA). Illumina 250 bp paired-end (250 PE) sequencing was undertaken on an Illumina MiSeq platform using MiSeq Reagent Kits v2.0 (500 cycles). The MiSeq was set to generate only FASTQ files with no post-run analysis. Samples were automatically demultiplexed by the MiSeq, based on the supplied index combinations.

### 2.9. Bioinformatic analysis

All loci were analysed separately using the same method as described in Chapter 3, Section 2.4. Briefly, amplicon sequence variants (ASVs) were identified for each locus using *dada2 v1.30.0* in RStudio. Filtering was employed to remove reads containing unresolved nucleotides as well as reads exceeding the expected error number and size range, *filterAndTrim* (run parameters: *maxN* = 0, *maxEE* = c(2,2), *truncQ* = 2, *minLen* = 50, *rm.phix* = TRUE, *matchIDs* = TRUE). This filtered dataset was then used for error training, *learnErrors(),* and error correction (denoising) of the dataset. The paired reads were merged, and chimeric sequences were identified before removal *removeBimeraDenovo().* A final table was produced for all the ASVs identified, along with their frequencies within the dataset.

### 2.10. Statistical analysis

All data were analysed and visualised with R Studio and publicly available packages: *tidyverse*, *drc*, *irr*, *tidymodels*, *eggCounts*, *bayescount*, and *ggplot2*. The EC_50_ (effective concentration for 50 % inhibition) and EC_95_ values for each EHT were calculated using the *drm()* function of the *drc* package using the LL.4 model (Ritz and Streibig, 2005). The *EDcomp()* function compares effective doses derived from dose-response curves and reports p-values reflecting the statistical significance of the differences between groups (p ≤ 0.05 considered significant). The resistance ratio was calculated as the EC_x_ value of the sample divided by the ECx of the susceptible isolate (from Farm FECRT_3).

For all FEC datasets, statistical analysis was performed to calculate the faecal egg count reduction (FECR) using *eggCounts* v2.4 to estimate the FECR with 90 % and 95 % confidence intervals (CI), and *bayescount* v0.9.99-9 to calculate the 90 % CI. The output of the *eggCounts* and *bayescount* packages were interpreted based on the original FECRT guidelines described by Coles et al., 1992 and the revised guidelines described by Kaplan et al., 2023. The two guidelines are summarised in Table 1. To quantitatively compare the agreement between statistical models and the original and revised FECRT guidelines, weighted Kohen’s κ coefficients were calculated using *kappa2(weights = “quadratic”)* (Normal < Inconclusive < Low resistant < Resistant) and interpreted as described by McHugh, 2012.

**Table 1.**
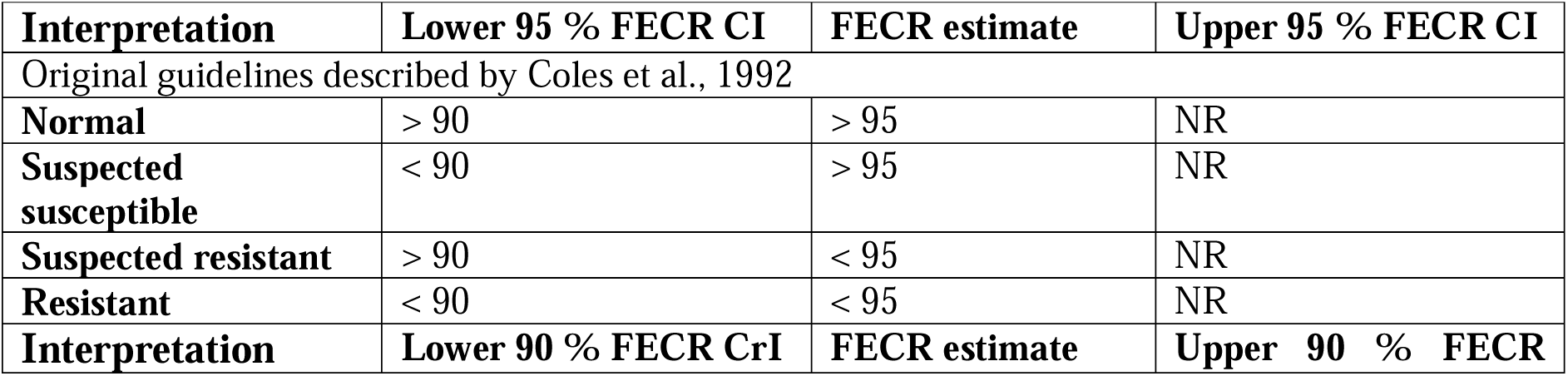
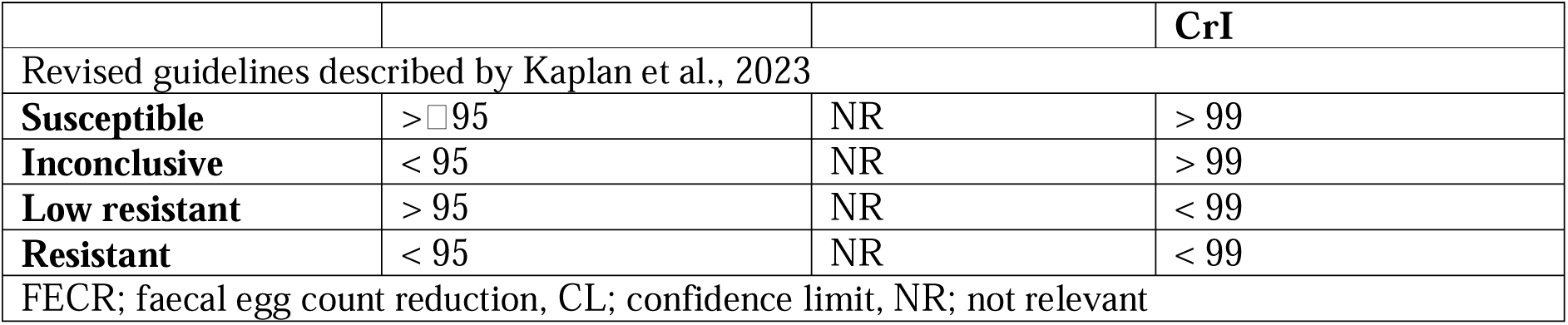
Comparison of the classification criteria for the faecal egg count reduction test.

## 3. Results

### 3.1. Farms included in the study and their parasite management

The basic farm demographics and anthelmintic use information are summarised in Table 2. Fourteen farms participated in this study: four farms participated in the FECRT, and ten organic farms participated in the egg hatch test. All farms were operated as commercial dairy farms, with a majority also engaged in dairy-beef production (13/14), and half (7/14) also had sheep. The groups of FGS calves ranged from 29 to 62 individuals, all of which were spring-born and aged between 4 and 7 months at the time of sampling, with 71 % to 100 % being female. All non-organic farms were reported to have exclusively used macrocyclic lactone anthelmintics during the previous seven years, while organic farms exclusively used BZ products, and no farms had reported using levamisole. All participating farms reported only administering anthelmintics at the group level. All non-organic farms reported an average of two anthelmintic treatments per year, broadly described as one mid-season treatment while at pasture and one at housing. Of the organic farms, 6/10 averaged one group treatment per year, while four reported fewer than one group treatment per year. There were three broad categories of treatment regimens employed by the farms in the study, which we define as follows:

- Neo-suppressive: treatment to limit the establishment of a parasitic infection and minimise pasture larval contamination
- Prophylactic: treatment of an at-risk group in anticipation of clinical or production-limiting parasitism based on previous management experience, but without the use of diagnostic indicators
- Test-and-treat: treatment based on an FEC that may be production-limiting

**Table 2.**
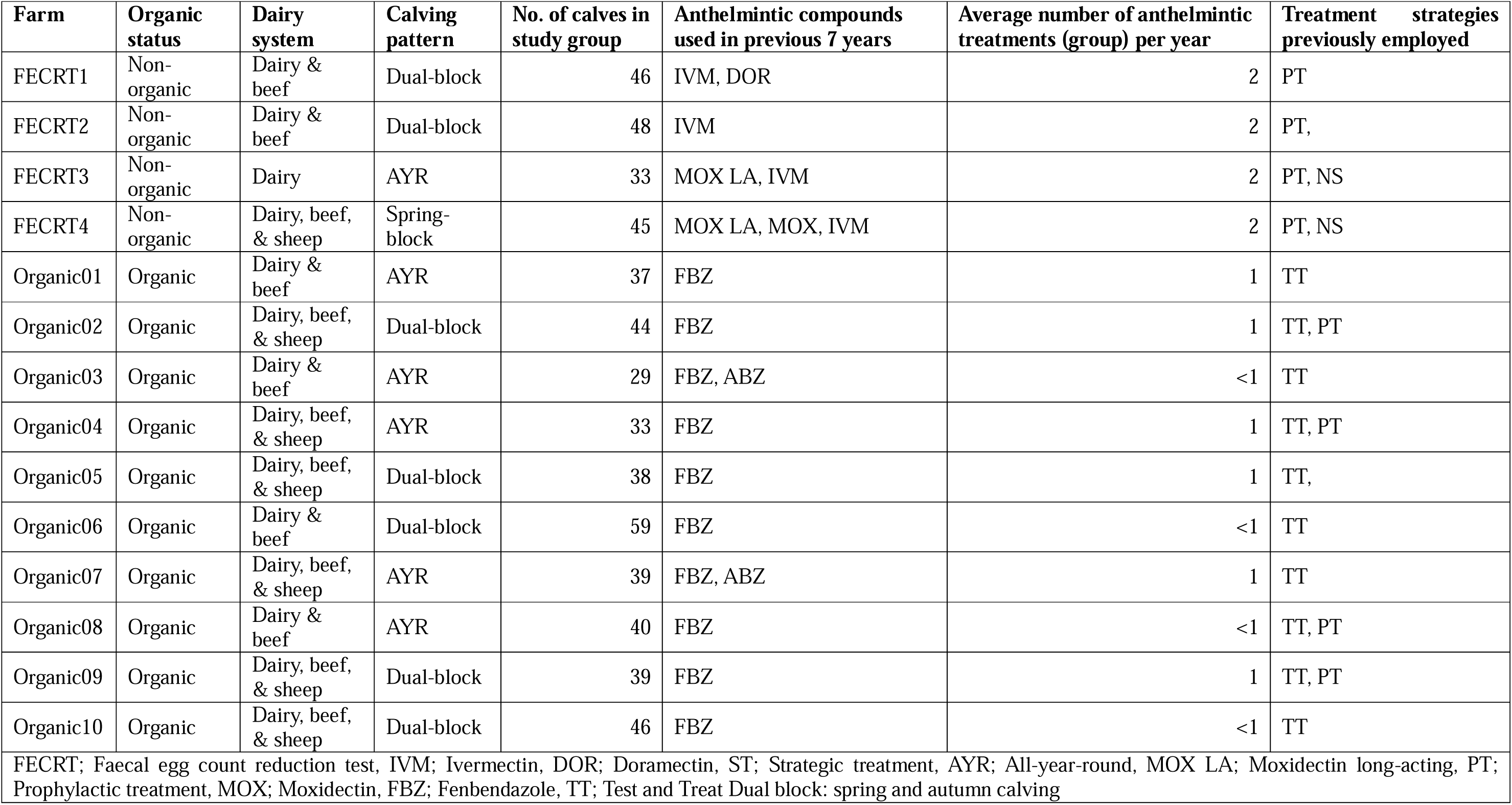
Farm demographics.

| **Farm** | **Organic status** | **Dairy system** | **Calving pattern** | **No. of calves in study group** | **Anthelmintic compounds used in previous 7 years** | **Average number of anthelmintic treatments (group) per year** | **Treatment strategies previously employed** |
| --- | --- | --- | --- | --- | --- | --- | --- |
| FECRT1 | Non-organic | Dairy & beef | Dual-block | 46 | IVM, DOR | 2 | PT |
| FECRT2 | Non-organic | Dairy & beef | Dual-block | 48 | IVM | 2 | PT, |
| FECRT3 | Non-organic | Dairy | AYR | 33 | MOX LA, IVM | 2 | PT, NS |
| FECRT4 | Non-organic | Dairy, beef, & sheep | Spring-block | 45 | MOX LA, MOX, IVM | 2 | PT, NS |
| Organic01 | Organic | Dairy & beef | AYR | 37 | FBZ | 1 | TT |
| Organic02 | Organic | Dairy, beef, & sheep | Dual-block | 44 | FBZ | 1 | TT, PT |
| Organic03 | Organic | Dairy & beef | AYR | 29 | FBZ, ABZ | <1 | TT |
| Organic04 | Organic | Dairy, beef, & sheep | AYR | 33 | FBZ | 1 | TT, PT |
| Organic05 | Organic | Dairy, beef, & sheep | Dual-block | 38 | FBZ | 1 | TT, |
| Organic06 | Organic | Dairy & beef | Dual-block | 59 | FBZ | <1 | TT |
| Organic07 | Organic | Dairy, beef, & sheep | AYR | 39 | FBZ, ABZ | 1 | TT |
| Organic08 | Organic | Dairy & beef | AYR | 40 | FBZ | <1 | TT, PT |
| Organic09 | Organic | Dairy, beef, & sheep | Dual-block | 39 | FBZ | 1 | TT, PT |
| Organic10 | Organic | Dairy, beef, & sheep | Dual-block | 46 | FBZ | <1 | TT |

FECRT; Faecal egg count reduction test, IVM; Ivermectin, DOR; Doramectin, ST; Strategic treatment, AYR; All-year-round, MOX LA; Moxidectin long-acting, PT; Prophylactic treatment, MOX; Moxidectin, FBZ; Fenbendazole, TT; Test and Treat Dual block: spring and autumn calving

All non-organic farms employed a prophylactic treatment regimen; while those that had also administered a moxidectin long-acting injectable also employed a neo-suppressive regimen. All organic farms used a test-and-treat approach using FECs, while 4/10 organic farms also employed a prophylactic treatment regimen, treating animals in anticipation of a significant parasite challenge and/or burden. Effective quarantine was not routinely practised on any farm in this study: five farms stated that they gave quarantine treatments to bought-in cattle, and all stated that they do not routinely treat every animal.

### 3.2. Faecal egg count reduction test

To meet the required minimum mean eggs counted required by the revised guidelines, one FEC per sample was needed to calculate and assign resistance status to the entire strongyle and *Os. ostertagi* communities. In comparison, three FEC per sample were required to obtain enough eggs to assign the resistance status to the *C. oncophora* populations of all farms. When the mean FEC of a sample was between 0 and 1 EPG, this was always rounded up to 1 EPG.

#### 3.2.1 Non-modelled data

After treatment with any anthelmintic drug, FECs were significantly reduced for all populations (Figs. 1-4), on all farms, regardless of the anthelmintic product used; the expected FECR is 99 % (Kaplan et al., 2023). The paired pre- and post-treatment faecal egg counts are depicted as violin plots in the first column of Figs 1-4, for all treatments, the pre-treatment data are over-dispersed, with significant variance in FEC. The paired FECR for each individual is shown in the second column, and a substantial difference in FECR was observed between *Os. ostertagi* and *C. oncophora* in each treatment. Species composition of each strongyle population was determined by multiplex PCR, revealing six species from five genera in the study. In every pretreatment population, *Os. ostertagi* was the most prevalent species, with *C. oncophora* always the second most prevalent species. In all post-treatment populations, only two species were observed: *Os. ostertagi* and *C. oncophora*. After FBZ treatment, the prevalence of *Os. ostertagi* consistently increased, whereas after MOX treatment, the prevalence of *C. oncophora* was consistently increased. In six of the eight IVM treatment populations, *C. oncophora* prevalence increased, while in the Farm FECRT_1 and FECRT_4 populations, *Os. ostertagi* was observed to increase post-treatment.

**Figure 1.**
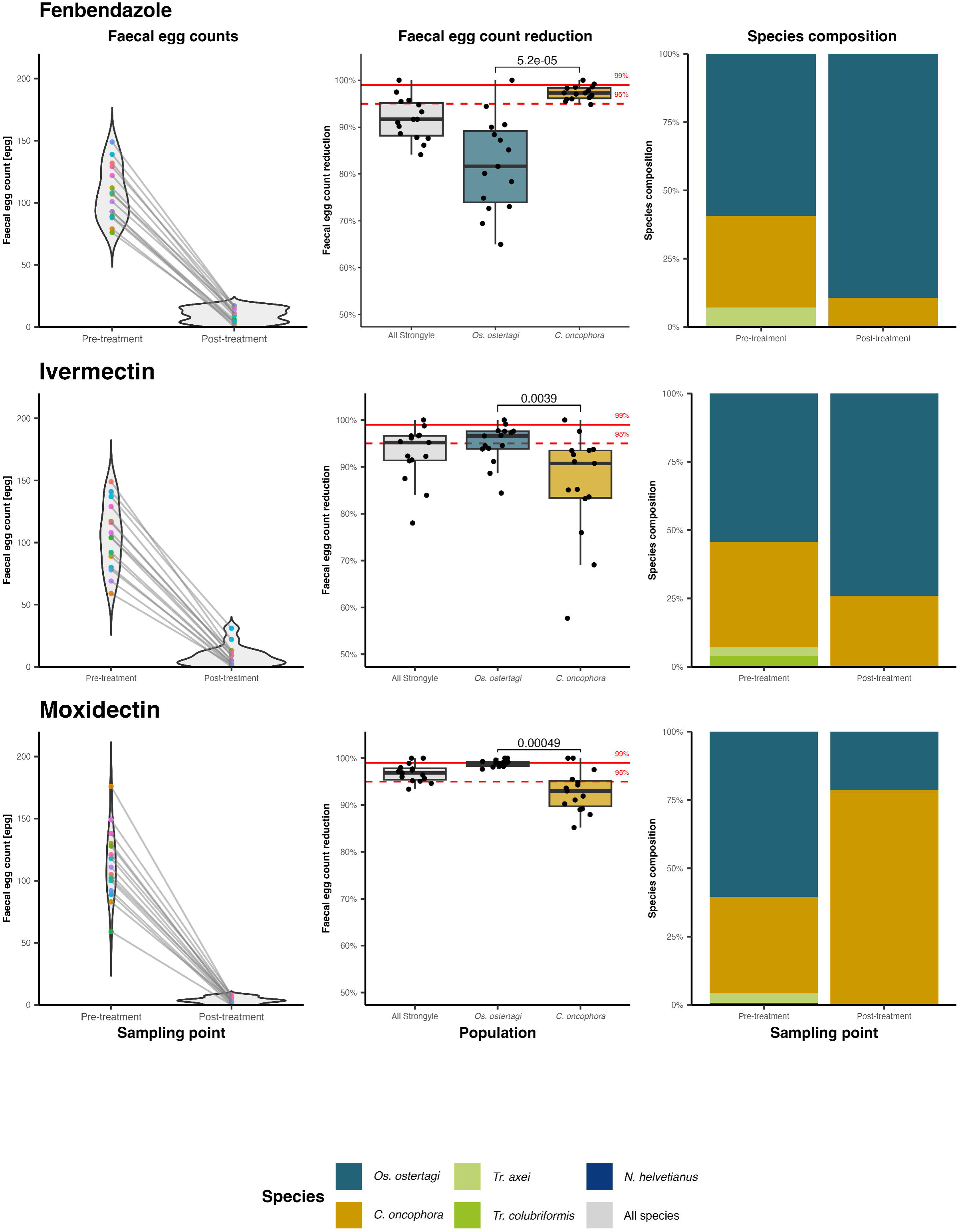
FECRT_1: paired faecal egg count reduction, anthelmintic efficacy and relative species abundance of gastrointestinal nematode communities, pre- and post-treatment. Species identity was assigned by ITS-2 rDNA multiplex PCR of a pool of L3 larvae harvested from coprocultures of each cohort and time point. A minimum of 94 L3 were identified per pooled coproculture. The violin plots with paired points represent the probability and distribution of the strongyle-type faecal egg counts (FECs) pre- and post-treatment. Faecal egg counts were conducted using a modified salt flotation technique with a sensitivity of epg 1. The boxplots represent the faecal egg count reduction estimates for each individual, based on the FEC and interpolated species compositions.

**Figure 2.**
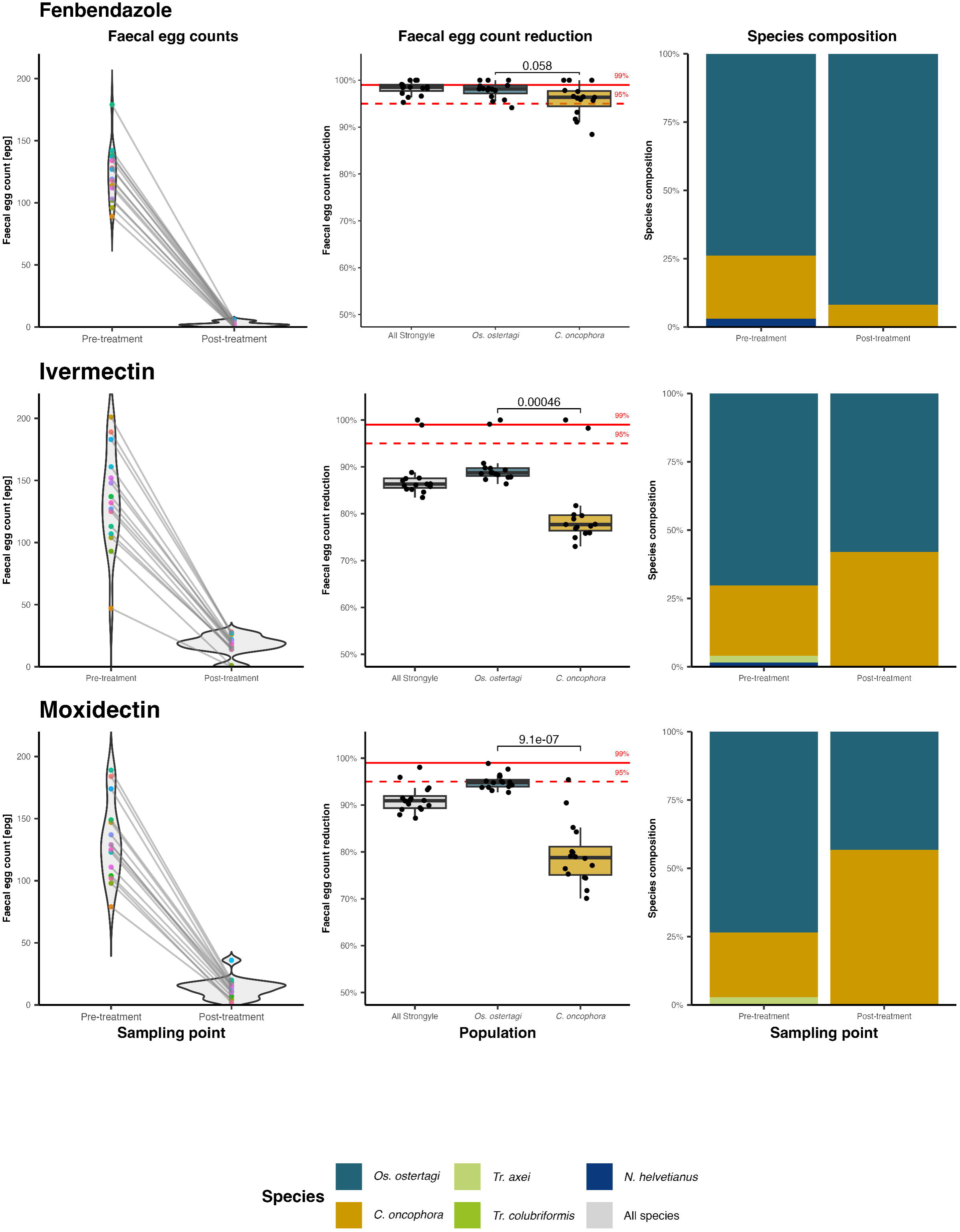
FECRT_2: paired faecal egg count reduction, anthelmintic efficacy and relative species abundance of gastrointestinal nematode communities, pre- and post-treatment. Species identity was assigned by ITS-2 rDNA multiplex PCR of a pool of L3 larvae harvested from coprocultures of each cohort and time point. A minimum of 94 L3 were identified per pooled coproculture. The violin plots with paired points represent the probability and distribution of the strongyle-type faecal egg counts (FECs) pre- and post-treatment. Faecal egg counts were conducted using a modified salt flotation technique with a sensitivity of epg 1. The boxplots represent the faecal egg count reduction estimates for each individual, based on the FEC and interpolated species compositions.

**Figure 3.**
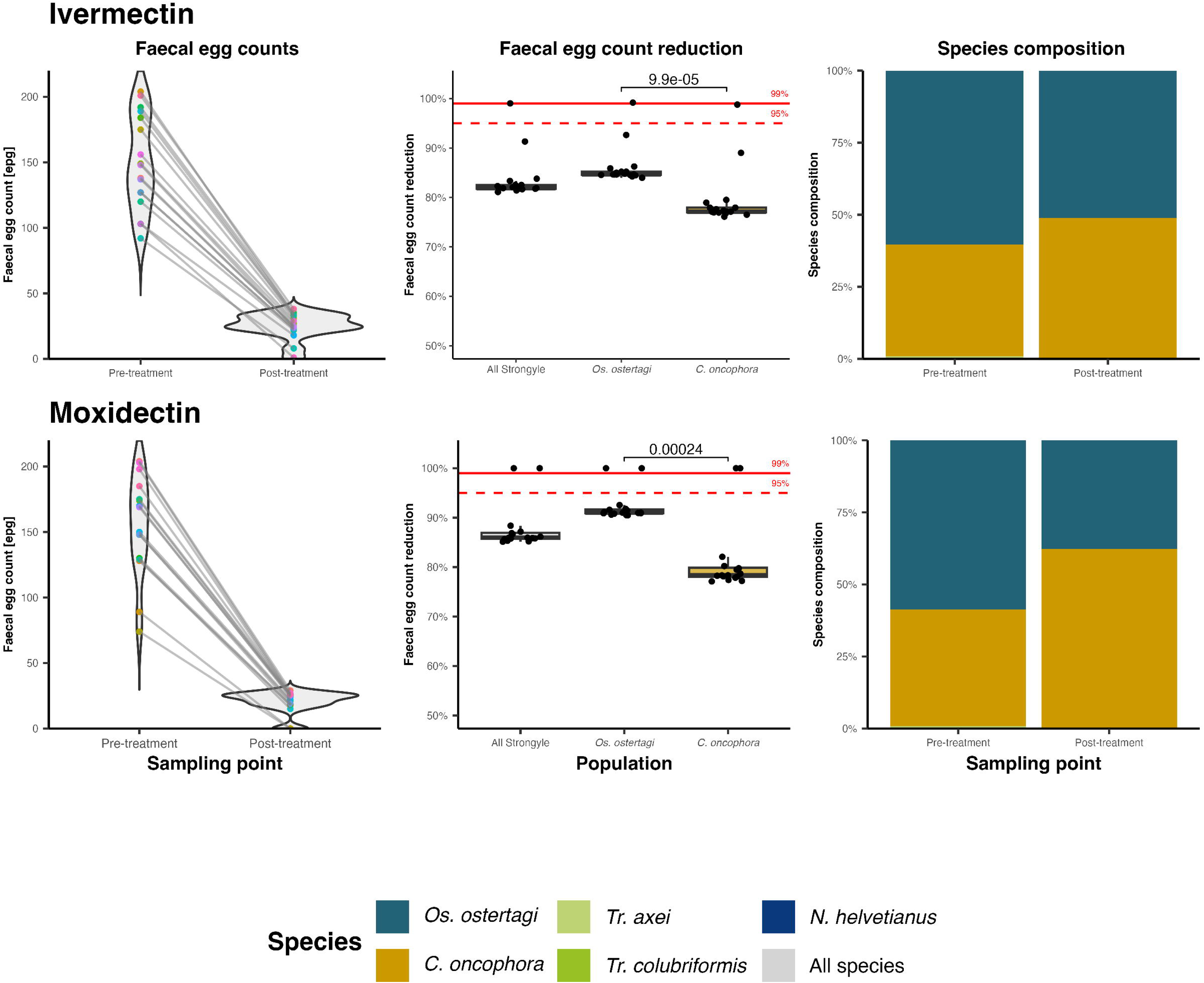
FECRT_3: paired faecal egg count reduction, anthelmintic efficacy and relative species abundance of gastrointestinal nematode communities, pre- and post-treatment. Species identity was assigned by ITS-2 rDNA multiplex PCR of a pool of L3 larvae harvested from coprocultures of each cohort and time point. A minimum of 94 L3 were identified per pooled coproculture. The violin plots with paired points represent the probability and distribution of the strongyle-type faecal egg counts (FECs) pre- and post-treatment. Faecal egg counts were conducted using a modified salt flotation technique with a sensitivity of epg 1. The boxplots represent the faecal egg count reduction estimates for each individual, based on the FEC and interpolated species compositions.

**Figure 4.**
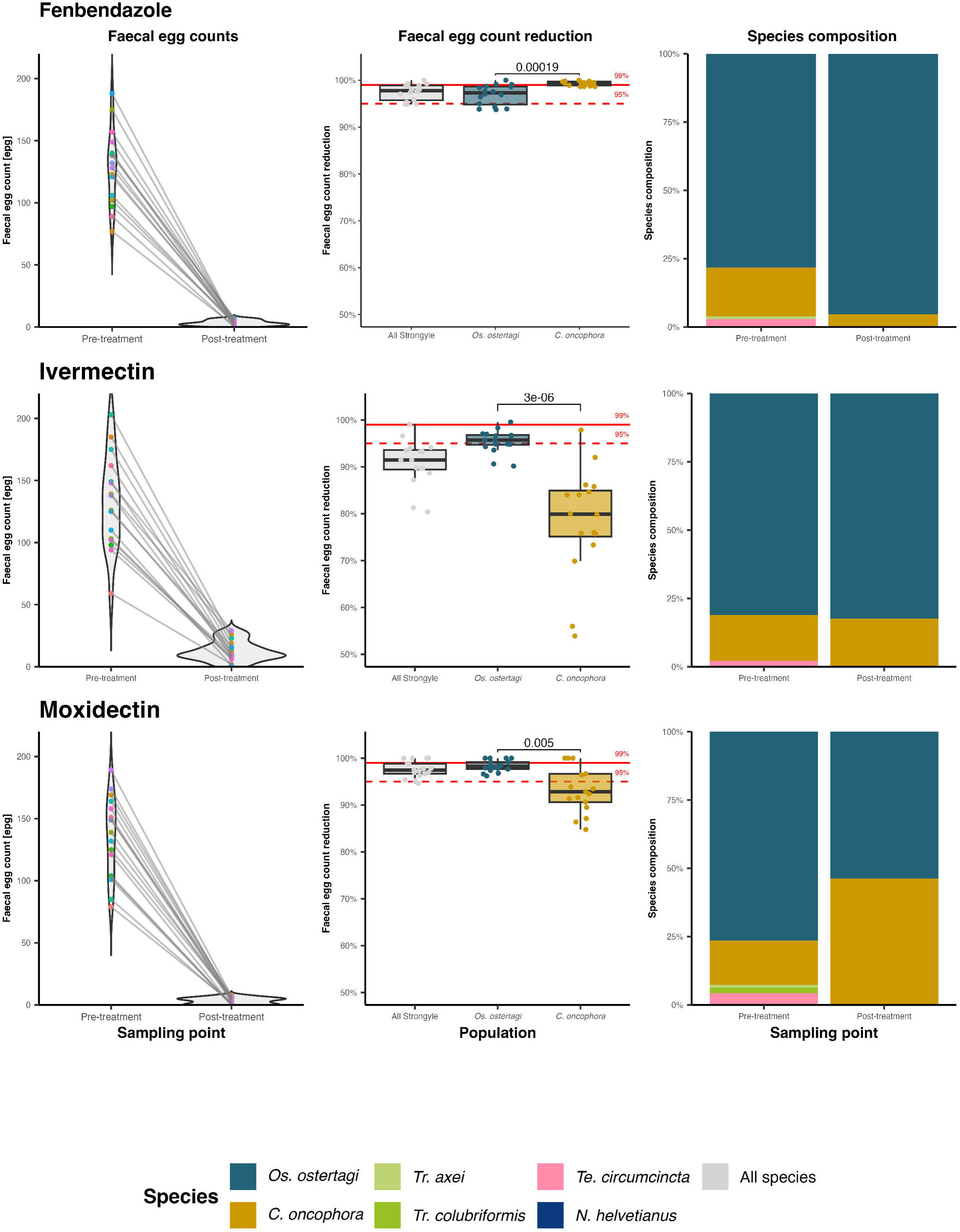
FECRT_4: paired faecal egg count reduction, anthelmintic efficacy and relative species abundance of gastrointestinal nematode communities, pre- and post-treatment. Species identity was assigned by ITS-2 rDNA multiplex PCR of a pool of L3 larvae harvested from coprocultures of each cohort and time point. A minimum of 94 L3 were identified per pooled coproculture. The violin plots with paired points represent the probability and distribution of the strongyle-type faecal egg counts (FECs) pre- and post-treatment. Faecal egg counts were conducted using a modified salt flotation technique with a sensitivity of epg 1. The boxplots represent the faecal egg count reduction estimates for each individual, based on the FEC and interpolated species compositions.

#### 3.2.1. Comparison and interpretation of the faecal egg count reduction test between statistical methods and guidelines

The interpretation of the FECRT using the previous WAAVP guidelines for the detection of anthelmintic resistance (Coles et al., 1992) was compared with the recently published revised guidelines (Kaplan et al., 2023). For this purpose, the categories for determining anthelmintic efficacy from the original guidelines - i.e., reduced, suspected resistant, suspected susceptible, and normal - were considered equivalent to those of resistant, low resistant, inconclusive, and susceptible, respectively, from the current revised guidelines. For all comparisons of guidelines and FECRT modelling packages, see Appendix File 5.

When assigning a resistance status to the entire strongyle population against FBZ using the revised guidelines, only one scenario resulted in a susceptible assignment (FECRT2/BZ/EC/RV), as shown in Fig. 5. Consistency between all FBZ scenarios was observed only in the status classified to FECRT_1, which classified the populations as resistant. In all IVM treatment scenarios in all farms, the strongyle populations were determined to be resistant. The MOX-treated populations of farms FECRT_2 AND FECRT_3 were consistently determined to be resistant in all scenarios, whereas the MOX populations of farms 1 and 4 were only classified as resistant in scenarios using the revised guidelines.

**Figure 5.**
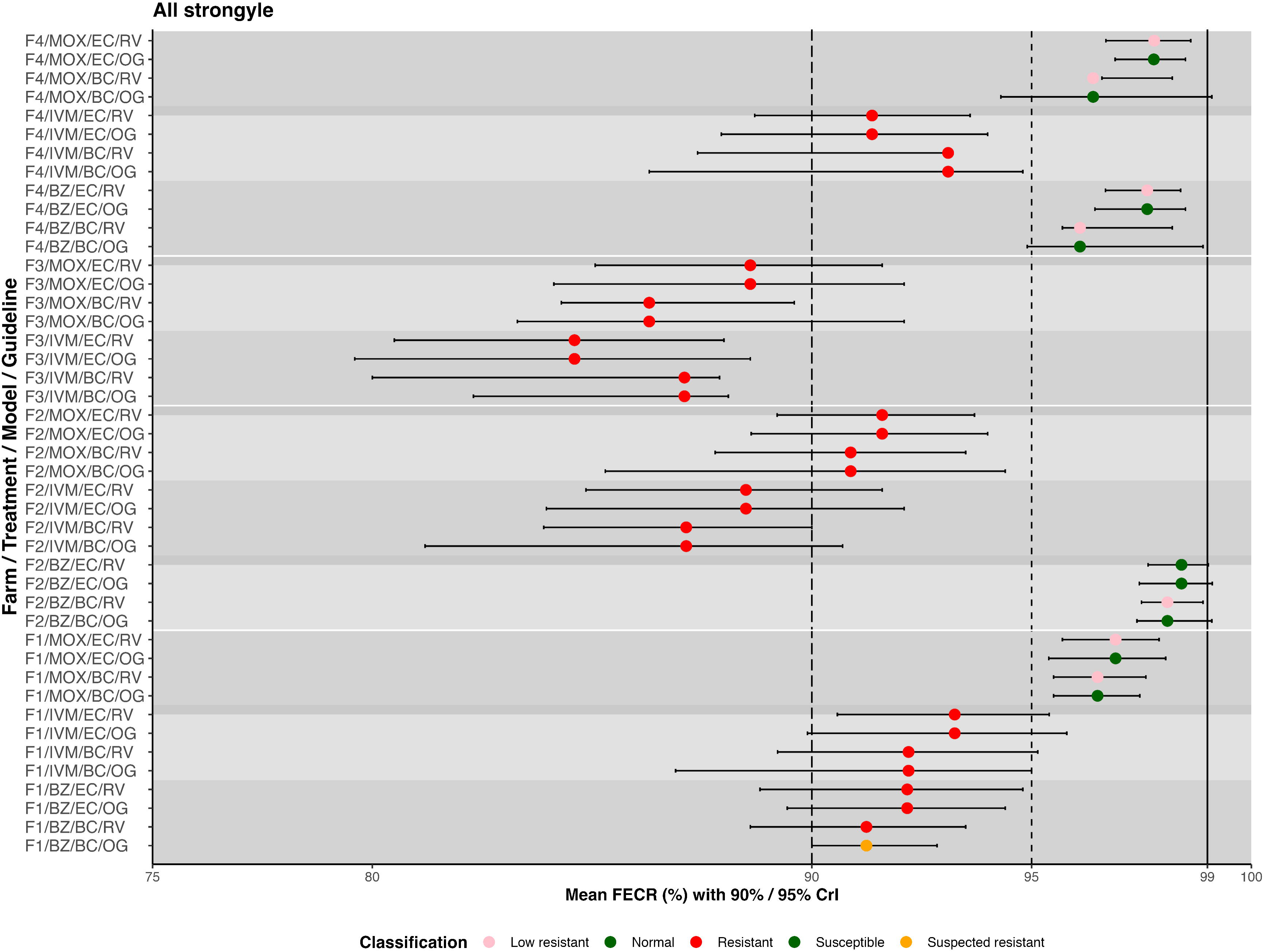
Comparison of the *eggCounts* and *bayescount* faecal egg count reduction estimates for the entire strongyle population. Faecal egg count reductions (FECR) with the credible intervals (CrIs) for anthelmintic treatment against the entire strongyle population. The CrIs were calculated using either *eggCounts* (EC) or *bayescount* (BC) models and interpreted based on either the revised guidelines (RV) for the faecal egg count reduction test (Kaplan et al., 2023) with corresponding 90 % CrIs, or on the original guidelines (OG) (Coles et al., 1992) with 95 % CrIs. Each point represents the mean FECR, with colour indicating the resistance status classified for the entire strongyle population: green, susceptible/normal; red, resistant; pink, low resistant/suspected resistant; orange, inconclusive/suspected susceptible.

Using the categories for each modelling / statistical method based on the revised guideline and treatment results, with the assignment of suspected resistance equal to low-resistance, both statistical methods agreed for 10/11 datasets (Table 3.). With one dataset classified as susceptible by *eggCounts* and classified as low resistance by *bayescount*. This resulted in a Cohen’s k value of 0.656, corresponding to substantial agreement.

**Table 3.**
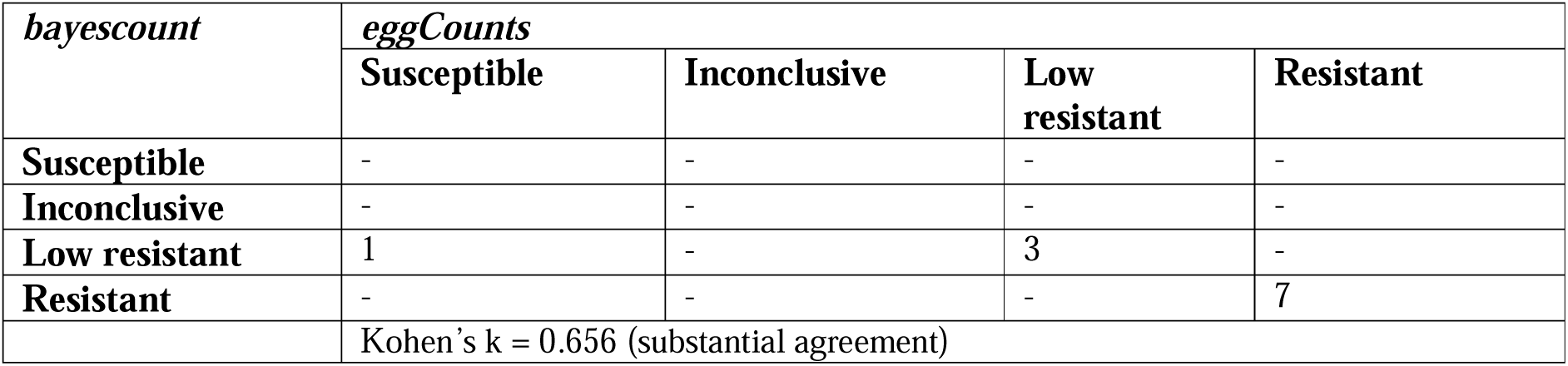
The inter-rater agreement between *eggCounts* and *bayescount* results based on the revised guidelines for the faecal egg count reduction test (Kaplan et al., 2023) for the entire strongyle population.

Comparing the interpretations of the FECRT results between the original and revised guidelines was conducted using only egg count data. The interpretation of results using both guidelines agreed for 8/11 datasets (Table 4.). Three datasets classified as low-resistant by the revised guidelines were classified as normal (susceptible) by the original guidelines. This results in the same level of agreement as between both modelling packages and the revised guidelines.

**Table 4.**
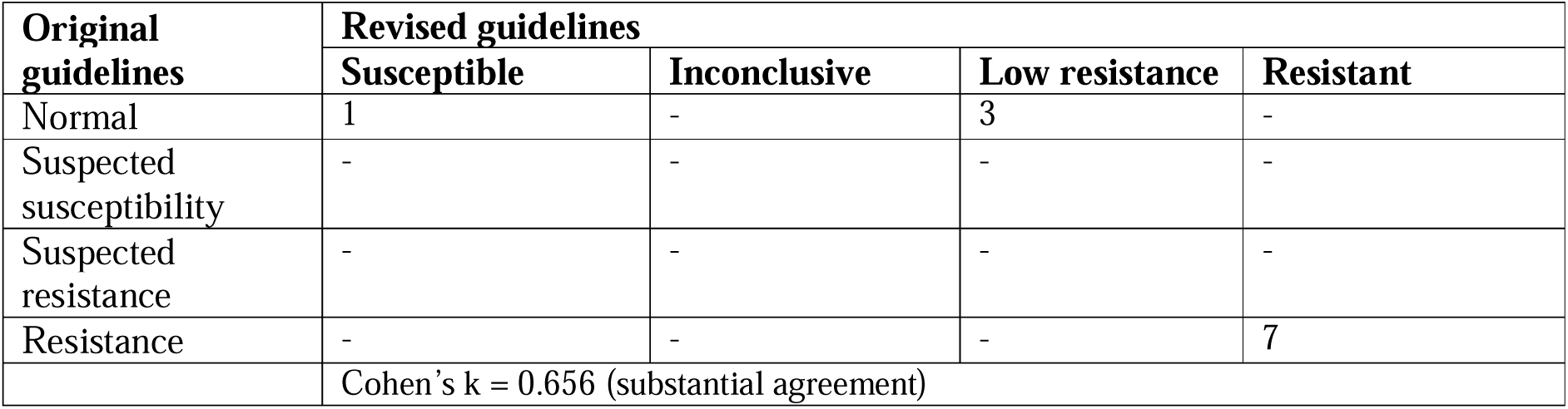
Inter-rater agreement between the original guidelines (Coles et al., 1992) and the revised guidelines (Kaplan et al., 2023) based on the faecal egg count reduction test for the entire strongyle population analysed using *eggCounts*.

When comparing the inter-rater agreement of the statistical models using the revised guidelines’ interpretations of the FECRT for the interpolated *Os. ostertagi* and *C. oncophora* populations. The *Os. ostertagi* populations were classified as the same statuses as the entire strongyle populations for all treatments (Appendix File X; Table X.). This resulted in a Cohen’s k value of 0.656, corresponding to substantial agreement. The inter-rater agreement for the *C. oncophora* populations was perfect, with a Cohen’s k value of 1, consistently assigning ten populations as resistant and one as low resistant.

Comparing the inter-rater agreement of the original and revised guidelines’ interpretation using the *eggCounts* model of the interpolated species datasets, there was fair agreement between the guidelines when assigning the resistant status to the *Os. ostertagi* populations with a Cohen’s k value of 0.333, where three populations were initially classified as normal by the original guidelines but were classified as low-resistant by the revised guidelines, and one population classified as normal was also classified as resistant. The inter-rater agreement for the *C. oncophora* populations was lower, with a Cohen’s k value of 0.19, consistently assigning resistance to eight populations; however, it also classified two resistant populations as normal by the original guidelines, as well as a low-resistant population and a normal population, corresponding to only slight agreement between the guidelines.

### 3.3. Egg hatch test dose-response

Sufficient numbers of eggs were collected from all organic farms and pre-treatment FECRT populations to conduct the egg hatch test. The mean number of eggs added to each well was 207, while the mean proportion of egg hatching in the control wells was 92.6 %. The dose-response curves for all populations (all strongyle, *Os. ostertagi*, and *C. oncophora*) are presented in Appendix File X and the effective concentrations are shown in Figure 6. The interpolated effective concentrations for *Os. ostertagi* ranged from EC_50_ (0.017 to 0.157 µg/ml) and EC_95_(0.045 to 1.293 µg/ml) and for *C. oncophora* , EC_50_ (0.025 to 0.111 µg/ml) and EC_95_ (0.416-0.859 µg/ml). The relative resistance ratio was also calculated for each species, with farm FECRT_3 used as the reference sensitive isolate. The RR ranged from 4.24 to 28.71, and 1.70 to 12.87 for *Os. ostertagi* and *C. oncophora,* respectively. For the specific EC_50_, EC_95_ and RR for each population, refer to Appendix File X.

**Figure 6.**
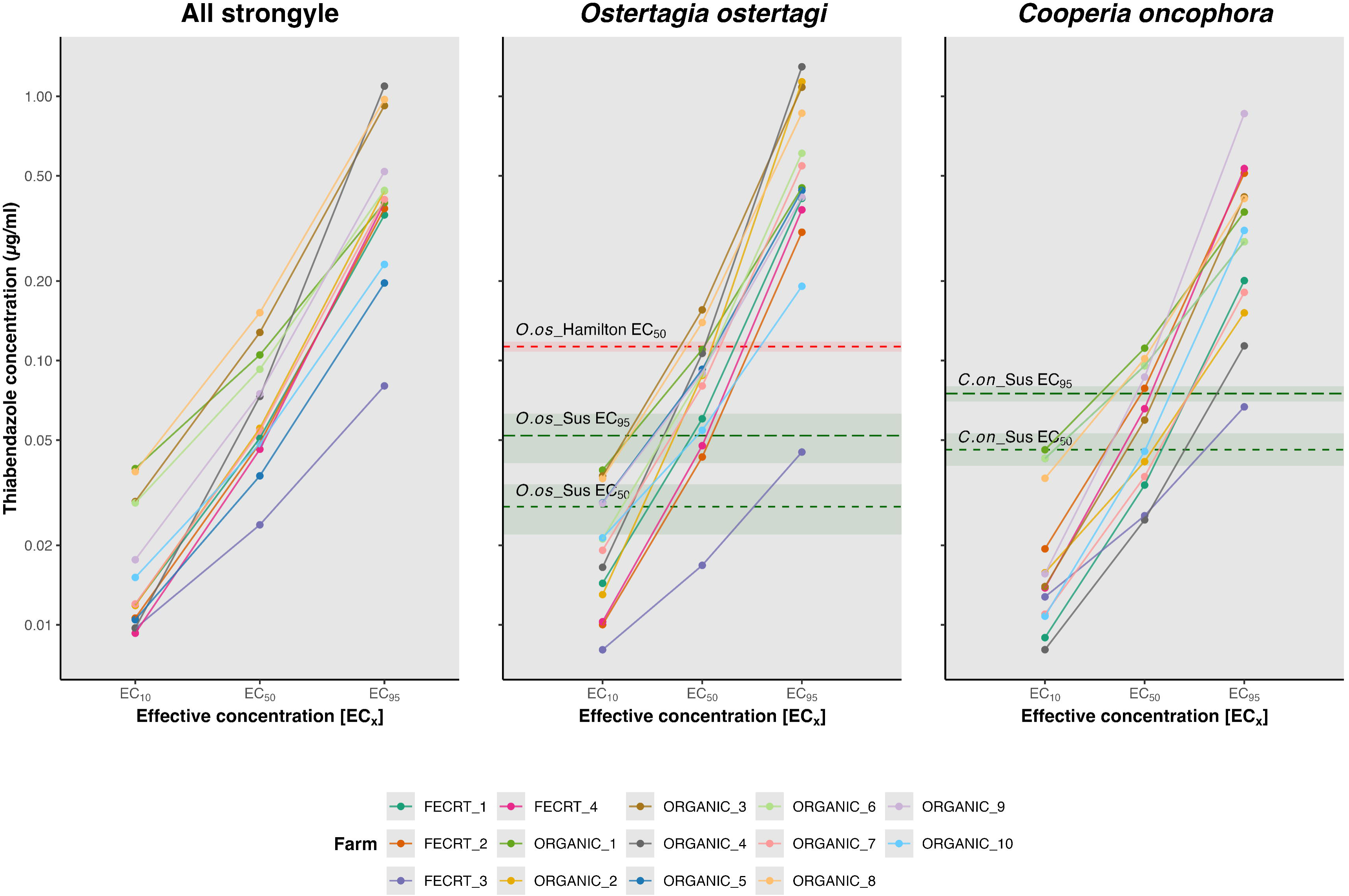
Nematode effective concentration estimates (μg/ml thiabendazole) for each strongyle and interpolated species population. The effective concentrations [E_C_] were estimated from the respective LL.4 dose-response curve model. The green dashed line represents the EC_50_ and EC_95_ of a fully susceptible *Ostertagia ostertagi* and *Cooperia oncophora* isolate, with the standard error of the mean. The red line represents the EC_50_ of a resistant *Os. ostertagi* population ± SEM.

### 3.4. Amplicon sequencing

Sufficient numbers of larvae were obtained from all treatment groups pre- (range 20,000 to 83,000) and post-treatment (range 2,300 to 5,100) to conduct mixed amplicon sequencing of all loci. Ten populations were subjected to the mixed amplicon sequencing marker panel based on the abundance of larvae, parasite management practices, and the likelihood of resistance being present, as determined by either or the FECRT or EHT. Four GIN populations from organic dairy farms and six paired pre- and post-treatment FECRT populations (FBZ, IVM, MOX) were selected.

#### 3.4.1. Nemabiome

We used ITS-2 nemabiome metabarcoding to determine the species composition of the GIN populations and identified 105 ASVs classified into eleven taxa, either at the genus (3.7 %) or species level (96.3 %). A minority of ASVs (n = 5) could not be classified to species but were identified as from the genera *Cooperia* (n = 3) and *Trichostrongylus* (n = 2). The most prevalent species identified were *Os. ostertagi* and *C. oncophora*, observed in every population (see Figure 7; Appendix 6). *Trichostrongylus axei* was observed on every farm but was absent in both the post-IVM and MOX populations, while *Tr. colubriformis* was observed on six farms. *Oesophagostomum* spp. were only observed in GIN populations from organic farms (n = 4), and *Haemonchus contortus* was observed on two farms. In terms of relative abundance, *Os. ostertagi* was the most abundant species in the majority of populations (8/10) and the most abundant species on all farms except farm FECRT_3. Comparing the changes in species abundance pre- and post-BZ-treatment, *Os. ostertagi* and *Teladorsagia circumcincta* increased from 53.1 % to 85.2 % and 1.5 % to 3.9 % respectively, while *C. oncophora* decreased from 29.7 % to 7.9 %. Comparing the effect of macrocyclic lactone products, *Os. Ostertagia* was observed to decrease from 67.2 % to 51.7 % and from 42.6 % to 28 %, post-IVM and MOX treatment, respectively, and *C. oncophora* was observed to increase from 21.3 % to 39.9 % and from 45.8 % to 71.1 %, respectively.

**Figure 7.**
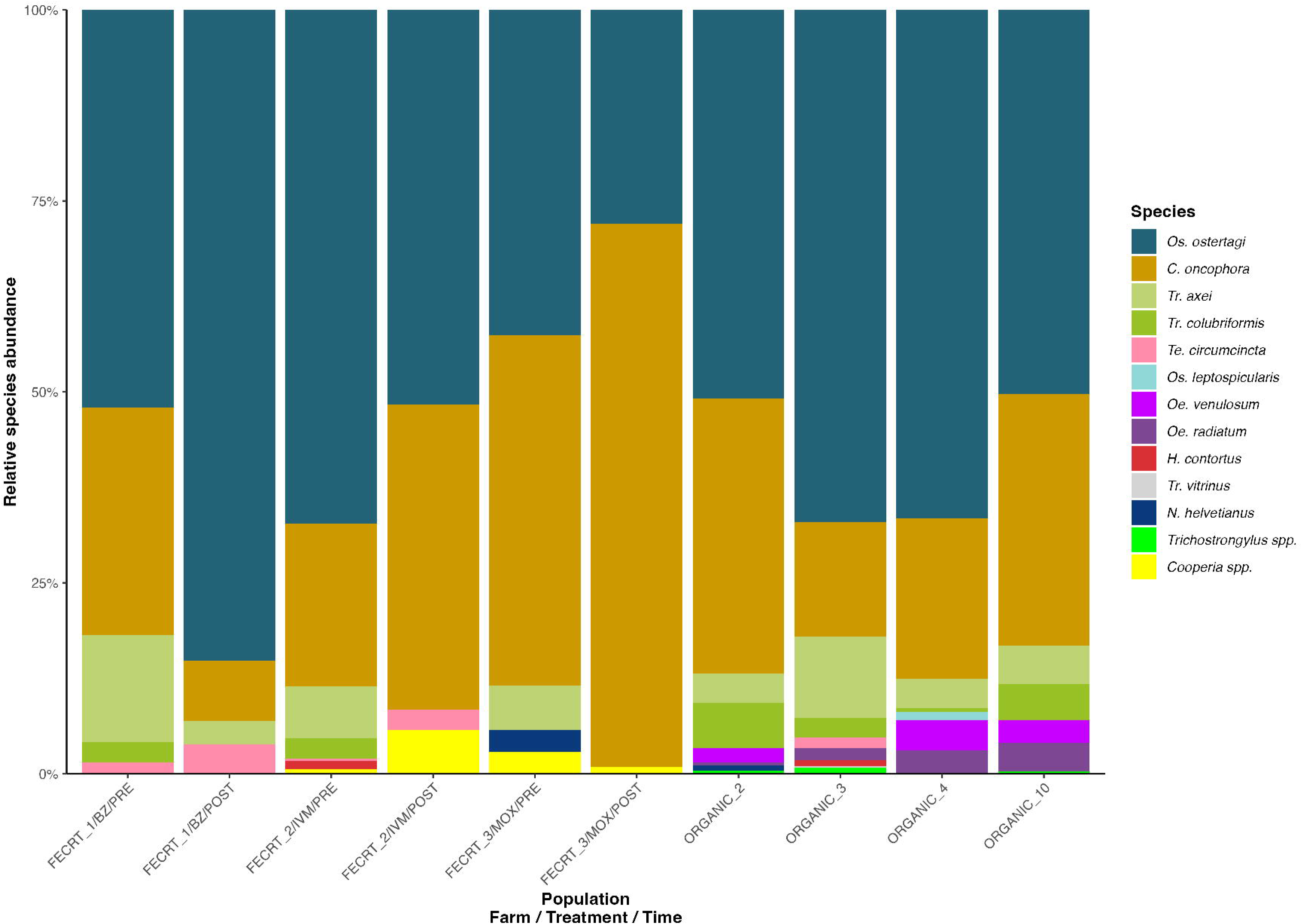
Gastrointestinal nematode composition determined by ITS-2 nemabiome metabarcoding. Relative species abundance of gastrointestinal nematode communities, determined by ITS-2 rDNA nemabiome metabarcoding, of pools of L_3_ harvested from individual coprocultures of pre- and post-faecal egg count reduction test populations and free-catch pasture samples from organic farms. A minority of ASVs could only be identified to the genus level: *Cooperia* spp. and *Trichostrongylus* spp. groups.

#### 3.4.2. Frequency of b-tubulin isotype-1 resistance-associated polymorphisms

Between 4,385 and 6,211 (mean = 5,381) sequence reads for the *b-tubulin isotype-1* loci were generated from each GIN population using the mixed amplicon sequencing panel and screened for BZ-resistance polymorphisms at codons 167, 198 and 200. Sequences mapped to *Os. ostertagi*, *Tr. axei*, *Tr. colubriformis*, *Tr. vitrinus*, *C. oncophora*, *Te. circumcincta*, *H. contortus* C*. c<u>urticei</u>*, *Os. leptospicularis*, and *Oesophagostomum* spp.. *Ostertagia ostertagi* and *C. oncophora B-tubulin* were identified in all samples (Figure 8). Benzimidazole resistance alleles were present to some degree in all strongyle populations from organic farms and were detected in seven of the ten sequenced populations. The majority of *Os. ostertagi* resistance alleles detected were F200Y (T<u>T</u>C > T**<u>A</u>**C), as well as two resistance alleles at codon 198, E198A (G<u>A</u>A > G**<u>C</u>**A), E198L (<u>GA</u>A > **<u>TT</u>**A). However, the F167Y (TTC > T**<u>A</u>**C) polymorphism was also present in organic farms 3 and 10 at very low frequencies of 1.3 % and 2.6 %, respectively. All *C. oncophora* resistance alleles were the F200Y variant, present at a mean frequency of 11.2 %, but showed high variability in frequency between farms, ranging from 2 % to 65.8 %. No non-synonymous polymorphisms were detected at codons 167 or 198. *Trichostrongylus* spp. (*Tr. axei*, *Tr. colubriformis*, *Tr. vitrinus*) alleles were detected on all farms but were absent in the post-macrocyclic lactone treatment populations. The F200Y polymorphism was detected at a mean frequency of 13.9 % in *Trichostrongylus* spp., while the E198L polymorphism was present at a highly variable frequency, with a mean of 11.5 %, ranging from 1.1 % to 43.4 %. Comparing the effect of fenbendazole treatment on allele frequency, the total frequency of resistance alleles for all strongyles increased from 12.7 % to 43.8 %, while for *Os. ostertagi*, *C. oncophora*, and *Trichostrongylus* spp., the frequencies increased from 20.8 to 35.6 %, 2 to 65.8 %, and 22.6 to 64.6 %, respectively.

**Figure 8.**
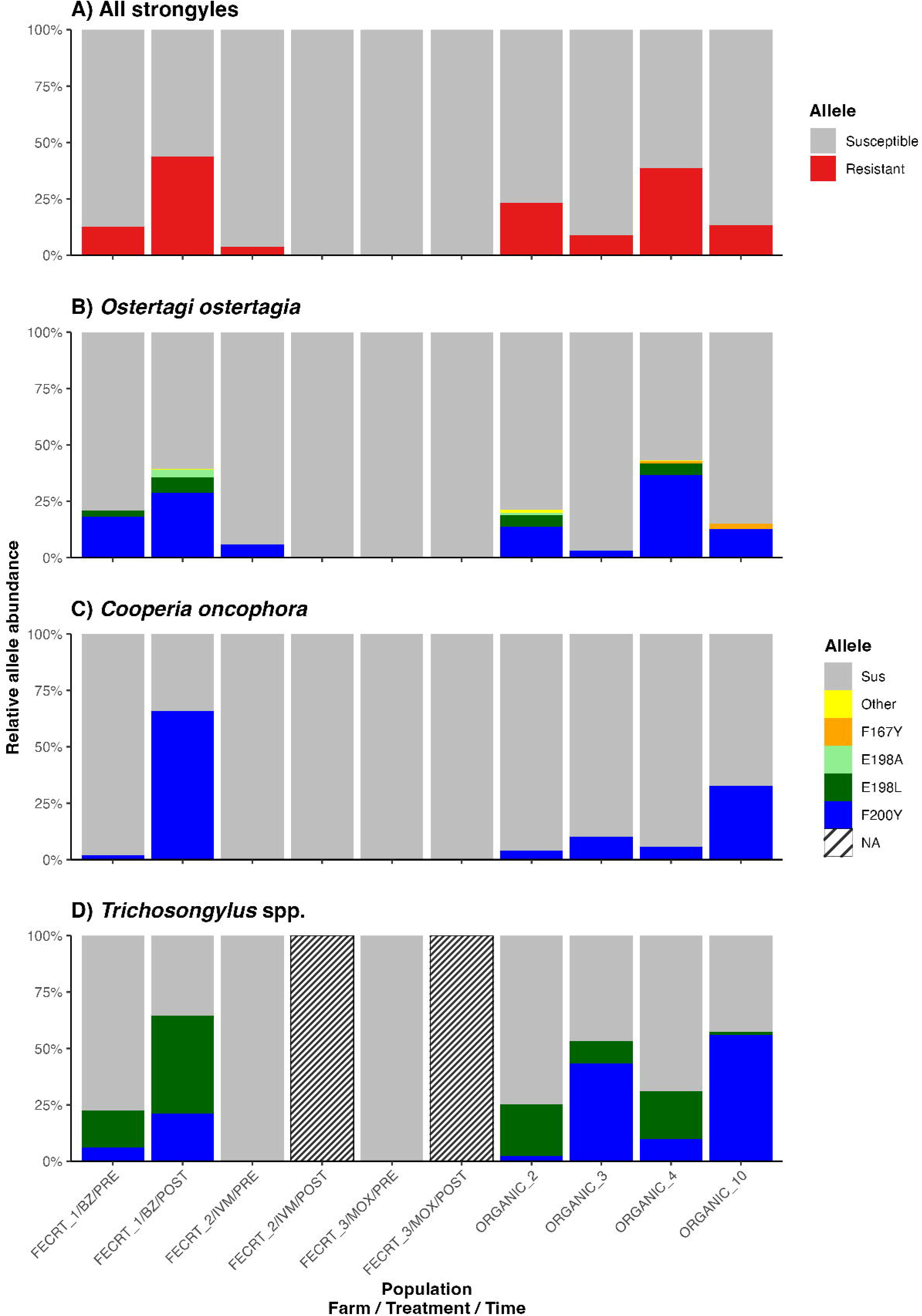
Frequency and prevalence of *b-tubulin isotype-1* gene resistance alleles. The relative proportions of the *b-tubulin isotype-1* gene resistance allele frequencies of 10 populations from seven different farms for all strongyles and three nematode species. (A) All strongyles; (B) *Ostertagia ostertagi*; (C) *Cooperia oncophora*; (D) *Trichostrongylus* spp. Susceptible alleles (F167, E198 and F200) are displayed in grey, while previously described resistance-associated polymorphisms (F167Y (T<u>T</u>C > T**<u>A</u>**C), E198A (G<u>A</u>A > G**<u>C</u>**A), E198L (<u>GA</u>A > **<u>TT</u>**A) and F200Y (T<u>T</u>C > T**<u>A</u>**C)) are displayed in red for all strongyles and as orange, green, light green and blue respectively for the resistance allele frequency per species. All other identified non-synonymous polymorphisms were grouped and are displayed in yellow. If no alleles were identified, this is represented as a diagonal stripe (NA).

**Frequency of *acr-8* and *b-tubulin isotype-2* resistance-associated polymorphisms**

Unambiguous *Os. ostertagi* S167T levamisole-resistant polymorphism (analogous to that of S168T in *H. contortus*) was detected in only one farm/population (ORGANIC_3) and was present at a low frequency of 1.1 %. However, the resistance allele was observed in two ASVs that cluster closely together phylogenetically. Fifty-seven ASVs were observed, generated from sequencing reads with depths ranging from 4,621 to 5,742 (mean = 5,297) per population. A second BZ resistance gene *b-tubulin isotype-2*, was screened for polymorphisms at codons 198 and 200; all ASVs generated were unambiguous for *Os. ostertagi,* and no resistance-conferring alleles were observed at the codon.

## 4. Discussion

The results of this study are consistent with the global trends of increasing resistance to all anthelmintic compounds and conclusively demonstrate that resistance to the main anthelmintic compounds exists in multiple nematode populations and species infecting Scottish dairy cattle. The control of GIN infections relies primarily on metaphylactic and therapeutic treatment with anthelmintics. In the UK and Europe, resistance in parasitic nematodes of cattle is currently not considered widespread and reported resistance levels have generally been low to moderate. The FECRT remains the most important field test for evaluating the susceptibility or resistance status of gastrointestinal nematode populations (Kotze et al., 2020). However, the original WAAVP guidelines for the FECRT (Coles et al., 1992) had not been updated in more than 30 years, until a recent revision by Kaplan et al., 2023. These revised guidelines not only update the recommendations for study design and reporting but also change the criteria (cut-offs) for identifying resistant populations. In addition, several different statistical approaches have been suggested to analyse FECRT data and to account for sources of variation (Denwood et al., 2023; Torgerson et al., 2014). It is essential to understand that the application of different guidelines and statistical approaches can yield varying interpretations of the same FECRT data. Ehnert et al., 2025, conducted a similar study and observed levels of agreement between the two statistical models consistent with this study, as well as low levels of agreement between the original and revised guidelines. In the current study, it should be emphasised that many of the interpolated species populations considered resistant by the revised guidelines were classified as susceptible when applying the original guidelines. This makes it difficult and unreliable to compare findings between studies using the original and revised guidelines, as it is likely that the original guidelines underestimate the prevalence of resistant strongyle populations.

The nemabiome analysis revealed that *Os. ostertagi* and *C. oncophora* were the most prevalent and abundant species in this study. This finding aligns with other UK and European studies, which have shown that these species dominate. The number of species identified in pre-treatment populations ranged from four to eight, totalling 11 species from six genera identified in the study. Despite the limitations of nemabiome analysis, cost, PCR bias, and egg shedding variation. It represents the best technique to characterise strongyle nematode communities. It is the only method that determines the species profile of strongyle nematode communities without making assumptions about their composition, while the volume of data generated enables the detection of rare species that are rarely studied. Correlation factors somewhat limit the impact of copy number variation between species; however, these limitations apply to all metabarcoding techniques. In most studies, read counts are used as a proxy for abundance.

After treatment with macrocyclic lactones, all farm populations showed an increase in the proportion of *Cooperia* spp. The generally low pathogenicity of this parasite could be a reason why resistance to the drug is rarely recognised by the farmers. In New Zealand, where most cases of GIN resistance to MLs have been reported, there are still no case reports of clinical parasitism due to this species (Jackson et al., 2006). Furthermore, the limited sensitivity of the FECRT, requiring at least 25% of a population to be resistant (Martin et al., 1989), in combination with the higher fecundity of *Cooperia* spp., likely underestimates efficacy against the less fecund *Os. ostertagi*. Additionally, the IVM concentration in the abomasal mucosa that *Os. ostertagi* experiences is higher than that of the intestine, the target site of *C. oncophora* (Lifschitz et al., 2000). This may be one reason why *Cooperia* spp. are the dose-limiting GIN. It was therefore expected that *Cooperia oncophora* would be the first species showing ML resistance in cattle (Coles, 2002).

In this study, the standardised egg hatch test protocol using thiabendazole was used to compare *in vitro* assay data, BZ-treatment FECRT data, and BZ-resistance allele frequency data. It is commonly accepted that the cut-off value for BZ-resistance is 0.1 μg TBZ/ml (Coles et al., 2006). The EC values reported in this study (EC_50_: 0.023 to 0.15 μg TBZ/ml, EC_95_: 0.08 to 1.07 μg TBZ/ml) were significantly higher than those obtained in a similar European study, where pre-treatment EC_50_ values of 0.027 to 0.038 μg TBZ/ml were observed for a mixed species population (Demeler et al., 2012). However, only one publication has given EC_50_ values for known susceptible *Os. ostertagi* and *C. oncophora* populations, which were reported at 0.022 to 0.034 μg and 0.04 to 0.052 μg, respectively. An EC_50_ value (0.108 to 0.118 μg) for a known resistant *Os. ostertagi* population (*O.o.*Hamilton, 2010) has been published (Demeler et al., 2013), but no efficacy values from an FECRT or controlled efficacy test have been reported. The extremely high EC_50_ and EC_95_ values for some populations are undoubtedly suggestive of a resistant population or subpopulation.

The mixed amplicon sequencing approach was successful in identifying known BZ-resistance alleles that were present in 0 to 43.8 % of all *b-tubulin isotype-1* reads and observed to significantly increase after fenbendazole treatment on farm FECRT_1. The F200Y variant was the most commonly observed resistance allele in *Os. ostertagi,* and the only resistance allele observed in *C. oncophora*. However, of the *Trichostrongylus* spp. resistance alleles, the E198L variant was the most commonly observed allele in 5/6 populations and was only more abundant on farm ORGANIC_10. The S167T variant is analogous to that of the S168T, which has been shown to be highly correlated with levamisole resistance in *H. contortus* (Antonopoulos et al., 2024), and was present at a very low frequency in two *Os. ostertagi* populations. The same analogous S167T variant in *Te. circumcincta* has been observed at high frequencies in one levamisole-resistant isolate (McIntyre et al., 2025). The presence of S167T in *Os. ostertagi* is a novel finding and has not been reported in this nematode species before. Although present at a low frequency, this resistance allele was detected in two ASVs, making it unlikely to have been caused by a PCR mismatch error.

## Conclusion

This study has examined the anthelmintic efficacy of the main anthelmintic compounds used to treat cattle in the UK, utilising both *in vivo* and *in vitro* assays, and highlighted the variance in anthelmintic efficacy between farms and strongyle species. The present study highlights the emergence of anthelmintic resistance to MLs and BZs in cattle nematodes, as well as the reliance of both conventionally and organically managed farms on these products. The variation in the efficacy of such treatments is evident, as well as the impact of differing statistical methodologies and diagnostic criteria on the assignment of resistance status. The revised FECRT guidelines complicate the comparability of results under previous guidelines and emphasise the need for methodological consistency. The higher-than-expected EC values for *Os. ostertagi* and *C. oncophora* are concerning and warrant further investigation, as there is a lack of empirical data and guidance on how to interpret such data and assign a resistance status. The variable frequency and high abundance of BZ-resistance alleles in some populations are noteworthy because, if the increasing trend for using BZ products against GIN infection in UK cattle continues, then selection for AR will likely rise. Consequently, continued and improved testing of efficacy to encourage remedial action would be very beneficial.

## Data Availability

Sequence data generated by this project were submitted to the SRA section of GenBank and are available under the SRA accession numbers within BioProject PRJNA. Additional information, data, and analysis code are available from GitHub (https://github.com/pau1campbe11/Characterising-AR-against-BZs-MLs-in-GINs.git )

## Acknowledgements

The authors thank all the participating farms and farm staff for their help and cooperation in this study. This work was funded by the James Herriot Scholarship (Glasgow University Vet Fund) (Paul Campbell). RL and JM were supported by a Wellcome Clinical Research Career Development Fellowship award to RL [216614/Z/19/Z] and a University of Glasgow Lord Kelvin Adam Smith Fellowship (LKAS) award to RL.

## Credit Authorship Contribution Statement

**Paul Campbell:** Conceptualisation, Data curation, Formal analysis, Investigation, Project administration, Writing – original draft, Writing – review & editing. **Jennifer McIntyre:** Conceptualisation, Formal analysis, Writing – review & editing. **Kerry O’Neill:** Investigation. **Andrew Forbes:** Conceptualisation, Formal analysis, Writing – review & editing. **Roz Laing:** Conceptualisation, Formal analysis, Supervision, Writing – review & editing. **Kathryn Ellis:** Conceptualisation, Formal analysis, Supervision, Writing – review & editing.

