## Supplementary for "Characterising anthelmintic resistance against benzimidazoles and macrocyclic lactones in gastrointestinal nematode populations of dairy cattle"

### Supplementary Information

### Supplementary file 1

**Table S1: Primer sequences for the ITS2 region used for gastrointestinal nematode identification, as developed by Bisset et al., 2014.**

| **Target species** | **Primer name** | **Sequence (5′–3’)** | **Melting temperature ($T_m$)** | **Expected amplicon size (bp)** |
|---|---|---|---|---|
| ITS2 Generic | ITS2GF | CACGAATTGCAGACGCTTAG | 54°C | 370-398 |
| | ITS2GR | GCTAAATGATATGCTTAAGTTCAGC | 54°C | |
| *Cooperia oncophora* | COONRV1 | CTATAACGGGATTTGTCAAAACAGA | 53°C | 173 |
| *Haemonchus spp.* | HACOFD3 | CATGTATGGCGACGATGTTCTT | 55°C | 90 |
| *Ostertagia ostertagi* | OSOSRV1 | CAATGTTAACGTCATGTTGCATTTCA | 55°C | 207 |
| *Trichostrongylus axei* | TRAXFD1 | CAAATATTGTGATAATTCCCATTTTAGTTT | 53°C | 236 |

The first four letters of each primer name indicate the target species: OSOS = *Ostertagia ostertagi*.
FD denotes forward primer
RV denotes reverse primer

### Supplementary file 2

**Table S2: Primer sequences for the *Ostertagia ostertagi* anthelmintic marker panel**

| **Target species** | **Target loci** | **Primer name** | **Sequence (5′–3’)** | **Melting temperature ($T_A$)** | **Cycles** | **Extension time (Seconds)** | **Expected amplicon size (bp)** | **Source** |
|---|---|---|---|---|---|---|---|---|
| Pan-nematode | ITS2 | NC1 | **TCGTCGGCAGCGTCAGATGTGTATAAGAGACAG**ACGTCTGGTTCAGGGTTGTT | | 40 | | | (Avramenko et al., 2015) |
| | | NC2 | **GTCTCGTGGGCTCGGAGATGTGTATAAGAGACAG**TTAGTTTCTTTTCCTCCGCT | | | | | |
| Pan-nematode | Beta-tubulin isotype-1 | Oos_tbb1_FW | **TCGTCGGCAGCGTCAGATGTGTATAAGAGACAG**ACGCACTCTTTGGGAGGAGG | | 40 | | | (Avramenko et al., 2019) |
| | | Con_tbb1_FW | **TCGTCGGCAGCGTCAGATGTGTATAAGAGACAG**TACGCATTCTCTTGGAGGAGG | | | | | |
| | | Oos_tbb1_RV | **GTCTCGTGGGCTCGGAGATGTGTATAAGAGACAG**TGTGAGTTTTAGTGTGCGGAAG | | | | | |
| | | Con_tbb1_RV | **GTCTCGTGGGCTCGGAGATGTGTATAAGAGACAG**TGTGAGCTTCAATGTGCGGAA | | | | | |
| | | Con/Hco_tbb1_FW | **TCGTCGGCAGCGTCAGATGTGTATAAGAGACAG**CGCATTCWCTTGGAGGAGG | | | | | |
| | | Con/Hco_tbb1_RV | **GTCTCGTGGGCTCGGAGATGTGTATAAGAGACAG**GTGAGYTTCAAWGTGCGGAAG | | | | | |

| | | | | | | | | |
|---|---|---|---|---|---|---|---|---|
| | | Tci/Tcol_tbb1_FW | **TCGTCGGCAGCGTCAGATGTGTATAAGAGACAG**CGCATTCYTTGGGAGGAGG | | | | | |
| | | Tci/Tcol_tbb1_RV | **GTCTCGTGGGCTCGGAGATGTGTATAAGAGACAG**GTGAGTTTYAAGGTGCGGAAG | | | | | |
| *Ostertagia ostertagi* | Beta-tubulin isotype-2 | Oos_tbb2_FW | **TCGTCGGCAGCGTCAGATGTGTATAAGAGACAG** | 51 | 40 | 15 | | This thesis, Chapter 3 |
| | | Oos_tbb2_RV | **GTCTCGTGGGCTCGGAGATGTGTATAAGAGACAG** | | | | | |
| *Ostertagia ostertagi* | cky-1 | Oos_cky1_FW | **TCGTCGGCAGCGTCAGATGTGTATAAGAGACAG** | 56 | 40 | 15 | | This thesis, Chapter 3 |
| | | Oos_cky1_RV | **GTCTCGTGGGCTCGGAGATGTGTATAAGAGACAG** | | | | | |
| *Ostertagia ostertagi* | acr-8 | Oos_acr8_FW | **TCGTCGGCAGCGTCAGATGTGTATAAGAGACAG** | 56 | 40 | 15 | | This thesis, Chapter 3 |
| | | Oos_acr8_RV | **GTCTCGTGGGCTCGGAGATGTGTATAAGAGACAG** | | | | | |
| *Ostertagia ostertagi* | pgp-9 | Oos_pgp9_FW | **TCGTCGGCAGCGTCAGATGTGTATAAGAGACAG** | 56 | 40 | 15 | | This thesis, Chapter 3 |
| | | Oos_pgp9_RV | **GTCTCGTGGGCTCGGAGATGTGTATAAGAGACAG** | | | | | |
| *Ostertagia ostertagi* | unc-29 | Oos_unc29_FW | **TCGTCGGCAGCGTCAGATGTGTATAAGAGACAG** | 56 | 40 | 15 | | This thesis, Chapter 3 |
| | | Oos_unc29_RV | **GTCTCGTGGGCTCGGAGATGTGTATAAGAGACAG** | | | | | |

| | | | | | | | | |
|---|---|---|---|---|---|---|---|---|
| *Ostertagia ostertagi* | unc-63 | Oos_unc63_FW | <u>**TCGTCGGCAGCGTCAGATGTGTATAAGAGACAG**</u> | 56 | 40 | 15 | | This thesis, Chapter 3 |
| | | Oos_unc63_RV | <u>**GTCTCGTGGGCTCGGAGATGTGTATAAGAGACAG**</u> | | | | | |
| *Ostertagia ostertagi* | chk | Oos_chk_FW | <u>**TCGTCGGCAGCGTCAGATGTGTATAAGAGACAG**</u> | 56 | 40 | 15 | | This thesis, Chapter 3 |
| | | Oos_chk_RV | <u>**GTCTCGTGGGCTCGGAGATGTGTATAAGAGACAG**</u> | | | | | |
| *Ostertagia ostertagi* | avr-14 | Oos_avr14_FW | **TCGTCGGCAGCGTCAGATGTGTATAAGAGACAG** | 56 | 40 | 15 | | This thesis, Chapter 3 |
| | | Oos_avr14_RV | **GTCTCGTGGGCTCGGAGATGTGTATAAGAGACAG** | | | | | |
| *Ostertagia ostertagi* | avr-15 | Oos_avr15_FW | **TCGTCGGCAGCGTCAGATGTGTATAAGAGACAG** | 56 | 40 | 30 | | This thesis, Chapter 3 |
| | | Oos_avr15_RV | **GTCTCGTGGGCTCGGAGATGTGTATAAGAGACAG** | | | | | |
| Ostertagia ostertagi | Neutral loci 1 | Oos_nloci_1_FW | **TCGTCGGCAGCGTCAGATGTGTATAAGAGACAG** | 56 | 40 | 15 | | This thesis, Chapter 3 |
| | | Oos_nloci_1_RV | **GTCTCGTGGGCTCGGAGATGTGTATAAGAGACAG** | | | | | |
| Ostertagia ostertagi | Neutral loci 2 | Oos_ nloci_2_FW | **TCGTCGGCAGCGTCAGATGTGTATAAGAGACAG** | 56 | 40 | 15 | | This thesis, Chapter 3 |
| | | Oos_ nloci_2_RV | **GTCTCGTGGGCTCGGAGATGTGTATAAGAGACAG** | | | | | |

| | | | | | | | | |
|---|---|---|---|---|---|---|---|---|
| Ostertagia ostertagi | Neutral loci 3 | Oos_ nloci_3_FW | **TCGTCGGCAGCGTCAGATGTGTATAAGAGACAG** | 56 | 40 | 15 | | This thesis, Chapter 3 |
| | | Oos_ nloci_3_RV | **GTCTCGTGGGC.AGATGTGTATAAGAGACAG** | | | | | |

The first three letters of each primer name indicate the target species: Oos = *Ostertagia ostertagi*.
FD denotes forward primer
RV denotes reverse primer

### Supplementary file 3

**Table S3: Individual population egg hatch test results**

The results of the egg hatch test performed with thiabendazole (TBZ) in 2023 on each study farm are presented at the extrapolated species and total strongyle level. Presented are the mean maximum effective concentrations (EC), $EC_{10}$, $EC_{50}$, and $EC_{95}$ values, with their 95 % confidence intervals (CI).

| Farm | Population | $EC_{10}$ µg TBZ/ml (CI) | $EC_{50}$ µg TBZ/ml (CI) | $EC_{95}$ µg TBZ/ml (CI) | RR $EC_{10}$ | RR $EC_{50}$ | RR $EC_{95}$ | AIC | Mean control hatch % |
|---|---|---|---|---|---|---|---|---|---|
| FECRT 1 | All strongyles | 0.0119337 (0.0012125) | 0.0509084 (0.0024112) | 0.3556748 (0.0423910) | 1.23113761 | 2.12894778 | 4.43518633 | 144.0472 | 92.7 |
| | *Os. ostertagi* | 0.0143691 (0.0030346) | 0.0602162 (0.0059686) | 0.4107894 (0.0993258) | 1.78542495 | 3.58378565 | 9.11692289 | 53.64525 | 92.8 |
| | *C. oncophora* | 0.0089455 (0.0018214) | 0.0338034 (0.0032040) | 0.2007497 (0.0481958) | 0.7011003 | 1.30589479 | 3.00535245 | 52.7031 | 91.3 |
| FECRT 2 | All strongyles | 0.0106009 (0.0013033) | 0.0487082 (0.0027461) | 0.3759052 (0.0552840) | 1.09363958 | 1.80552657 | 4.68745495 | 150.9229 | 91.2 |
| | *Os. ostertagi* | 0.0100143 (0.0018932) | 0.0431746 (0.0037479) | 0.3059468 (0.0716763) | 1.24432157 | 4.67221349 | 6.79008121 | 51.41523 | 93.9 |
| | *C. oncophora* | 0.0193927 (0.0046827) | 0.0785044 (0.0085789) | 0.5112810 (0.1385647) | 1.51989579 | 3.03278625 | 7.65420625 | 55.15238 | 89.9 |
| FECRT 3 | All strongyles | 0.00969323 (0.00073598) | 0.02391247 (0.00070135) | 0.08019388 (0.00664311) | NA* | NA* | NA* | 131.1158 | 96.4 |
| | *Os. ostertagi* | 0.0080480 (0.0011563) | 0.0168024 (0.0011492) | 0.0450579 (0.0081578) | NA* | NA* | NA* | 51.24016 | 97.6 |
| | *C. oncophora* | 0.01275923 (0.00112194) | 0.02588524 (0.00077946) | 0.06679739 (0.00679301) | NA* | NA* | NA* | 40.5817 | 96.2 |
| FECRT 4 | All strongyles | 0.0092948 (0.0013621) | 0.0461786 (0.0028377) | 0.3957303 (0.0760966) | 0.95889605 | 1.9311514 | 4.93466958 | 154.5357 | 92.5 |
| | *Os. ostertagi* | 0.0102716 (0.0030392) | 0.0476184 (0.0058641) | 0.3719197 (0.1512479) | 1.27629225 | 2.83402371 | 8.25426174 | 56.85585 | 93.8 |
| | *C. oncophora* | 0.0138007 (0.0021740) | 0.0657557 (0.0047907) | 0.5327762 (0.1002479) | 1.08162483 | 2.54027778 | 7.97600325 | 48.10137 | 91.9 |

| | | | | | | | | | |
|---|---|---|---|---|---|---|---|---|---|
| Organic 1 | All strongyles | 0.0390151 (0.0021380) | 0.1050033 (0.0025842) | 0.3957252 (0.0244986) | 4.02498445 | 4.3911524 | 4.93460598 | 123.4612 | 92.7 |
| | *Os. ostertagi* | 0.0385068 (0.0045677) | 0.1101367 (0.0057687) | 0.4503321 (0.0577333) | 4.78464215 | 6.55481955 | 9.99452038 | 46.29631 | 92.7 |
| | *C. oncophora* | 0.0459672 (0.0031150) | 0.1113913 (0.0034601) | 0.3647367 (0.0299381) | 3.60266254 | 4.30327476 | 5.46034359 | 39.56164 | 92.7 |
| Organic 2 | All strongyles | 0.0118376 (0.0010963) | 0.0554326 (0.0022466) | 0.4388187 (0.0503764) | 1.22122347 | 2.31814614 | 5.47197242 | 135.5862 | 92 |
| | *Os. ostertagi* | 0.0130175 (0.0022133) | 0.0878500 (0.0060138) | 1.1348755 (0.2236040) | 1.6174826 | 5.22841975 | 25.1870482 | 45.5948 | 91.3 |
| | *C. oncophora* | 0.0157589 (0.0017693) | 0.0414654 (0.0018623) | 0.1516112 (0.0222332) | 1.23509804 | 1.60189359 | 2.26971742 | 44.12307 | 91.3 |
| Organic 3 | All strongyles | 0.0292810 (0.0029930) | 0.1278455 (0.0057728) | 0.9214350 (0.1115102) | 3.0207681 | 5.34639458 | 11.4900913 | 143.9322 | 91.4 |
| | *Os. ostertagi* | 0.0365974 (0.0067762) | 0.1556091 (0.0122075) | 1.0823918 (0.2383759) | 4.54739066 | 9.26112341 | 24.0222425 | 51.28981 | 94 |
| | *C. oncophora* | 0.0139624 (0.0017768) | 0.0595655 (0.0034937) | 0.4161831 (0.0621494) | 1.09429801 | 2.30113764 | 6.23052937 | 45.36691 | 93.5 |
| Organic 4 | All strongyles | 0.0097278 (0.0018272) | 0.0731594 (0.0056526) | 1.0927365 (0.2320984) | 1.00356641 | 3.05946646 | 13.6261832 | 163.7521 | 92.4 |
| | *Os. ostertagi* | 0.0165157 (0.0069141) | 0.1064716 (0.0177297) | 1.2936106 (0.6024265) | 2.0521496 | 6.33669 | 28.7099621 | 61.46076 | 89.3 |
| | *C. oncophora* | 0.0080526 (0.0012904) | 0.0249623 (0.0016390) | 0.1136916 (0.0222276) | 0.63111959 | 0.96434493 | 1.70203656 | 49.38255 | 91.6 |
| Organic 5 | All strongyles | 0.01045391 (0.00077104) | 0.03662733 (0.00116865) | 0.19656662 (0.01791606) | 1.07847539 | 1.53172508 | 2.45114241 | 128.2538 | 92.6 |
| | *Os. ostertagi* | 0.0289504 (0.0027579) | 0.0926374 (0.0038906) | 0.4402493 (0.0503271) | 3.5972167 | 5.51334333 | 9.77074608 | 47.59801 | 92.9 |
| | *C. oncophora* | 0.0426220 (0.0019649) | 0.0955249 (0.0019842) | 0.2817013 (0.0164751) | 3.34048371 | 3.69032313 | 4.21725011 | 35.4871 | 92.9 |
| Organic 6 | All strongyles | 0.0289504 (0.0027579) | 0.0926374 (0.0038906) | 0.4402493 (0.0503271) | 2.98666182 | 3.87402054 | 5.48981169 | 144.7402 | 90.8 |
| | *Os. ostertagi* | 0.0211555 (0.0019548) | 0.0888857 (0.0035527) | 0.6084787 (0.0662158) | 2.62866551 | 5.29005975 | 13.5043733 | 39.98248 | 94.4 |
| | *C. oncophora* | 0.0426220 (0.0019649) | 0.0955249 (0.0019842) | 0.2817013 (0.0164751) | 3.34048371 | 3.69032313 | 4.21725011 | 33.51425 | 91.3 |
| Organic 7 | All strongyles | 0.0119998 (0.0011622) | 0.0540808 (0.0022209) | 0.4066997 (0.0454871) | 1.2379568 | 2.26161496 | 5.07145558 | 137.4504 | 94.2 |

| | | | | | | | | | |
|---|---|---|---|---|---|---|---|---|---|
| | *Os. ostertagi* | 0.0191599 (0.0033876) | 0.0801495 (0.0060025) | 0.5454631 (0.1120522) | 2.38070328 | 4.77012213 | 12.1058261 | 49.73166 | 93.3 |
| | *C. oncophora* | 0.0109699 (0.0017339) | 0.0363695 (0.0023311) | 0.1812554 (0.0359333) | 0.85976191 | 1.4050285 | 2.71351021 | 48.02988 | 92.7 |
| Organic 8 | All strongyles | 0.0379902 (0.0082538) | 0.1518750 (0.0127264) | 0.9726588 (0.2210851) | 3.91925086 | 6.35128868 | 12.1288408 | 174.2375 | 91.5 |
| | *Os. ostertagi* | 0.0357684 (0.0074248) | 0.1393750 (0.0112714) | 0.8624724 (0.1924796) | 4.4443837 | 9.0388873 | 19.1414247 | 51.37389 | 91.5 |
| | *C. oncophora* | 0.0358603 (0.0042376) | 0.1016091 (0.0052023) | 0.4102727 (0.0521597) | 2.81053794 | 5.38434258 | 6.14204687 | 45.6656 | 91.5 |
| Organic 9 | All strongyles | 0.0176462 (0.0016217) | 0.0748474 (0.0030827) | 0.5189282 (0.0584032) | 1.82046645 | 4.24920972 | 6.47092022 | 139.2358 | 92.9 |
| | *Os. ostertagi* | 0.0288277 (0.0023112) | 0.0900963 (0.0033897) | 0.4148617 (0.0410180) | 3.58197068 | 4.45456601 | 9.20730216 | 40.7812 | 94.3 |
| | *C. oncophora* | 0.0156165 (0.0021274) | 0.0865862 (0.0051030) | 0.8595706 (0.1422484) | 1.22393749 | 3.48060516 | 12.8683261 | 44.33899 | 91.9 |
| Organic 10 | All strongyles | 0.0151232 (0.0013673) | 0.0485001 (0.0019124) | 0.2311808 (0.0242190) | 1.56018169 | 2.02823464 | 2.8827736 | 140.2134 | 91.7 |
| | *Os. ostertagi* | 0.0213525 (0.0027061) | 0.0544768 (0.0028198) | 0.1911193 (0.0281091) | 2.65314364 | 3.24220349 | 4.24163798 | 46.68144 | 91,9 |
| | *C. oncophora* | 0.0107845 (0.0029378) | 0.0453442 (0.0057131) | 0.3107092 (0.1028604) | 0.84523126 | 1.7517396 | 4.65151707 | 57.15371 | 91.7 |

EC, Maximum effective concentration; TBZ, Thiabendazole; CI, Confidence interval; RR, Resistance ratio; AIC, Akaike Information Criterion; FECRT, faecal egg count reduction test

* Susceptible population used to calculate relative ratio

### Supplementary file 4

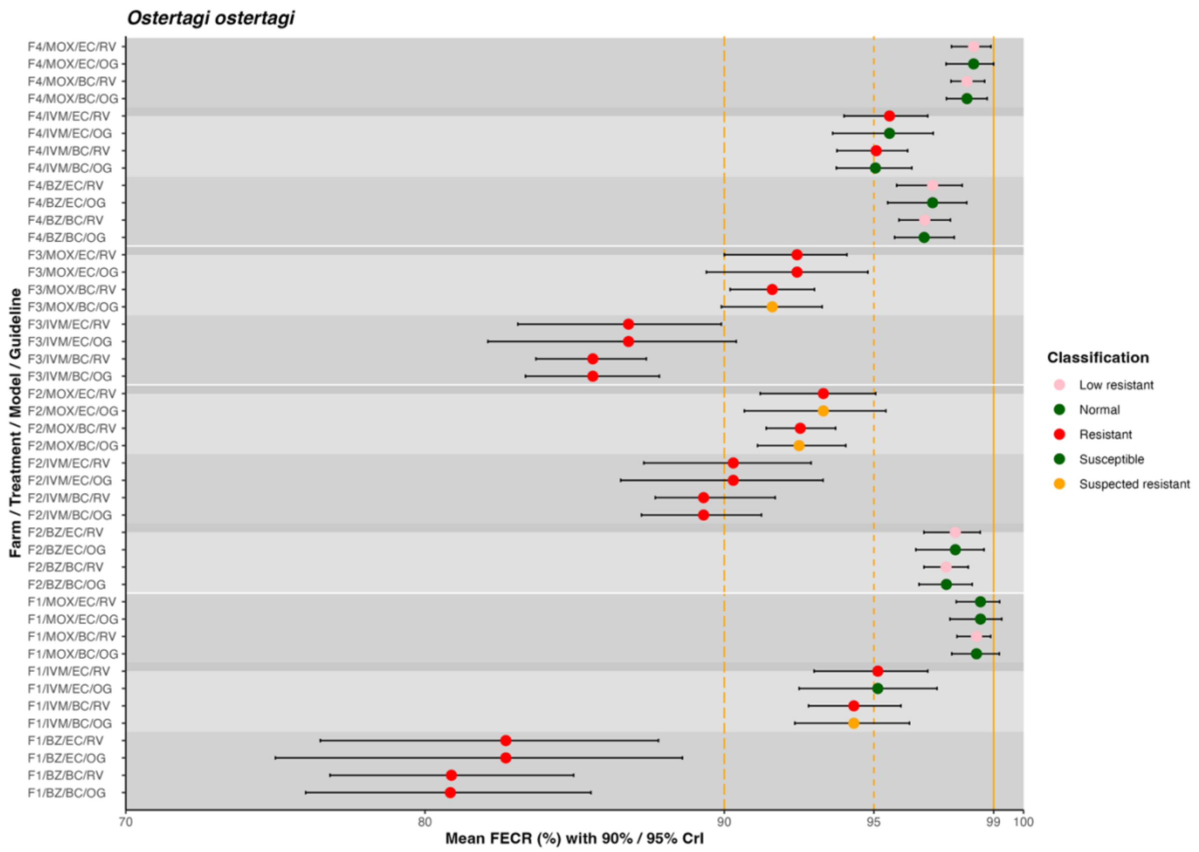


**Figure S1: Comparison of the *eggCounts* and *bayescount* faecal egg count reduction estimates for the interpolated *Ostertagia ostertagi* population**

Faecal egg count reductions (FECR) with the credible intervals (CrIs) for anthelmintic treatment against the interpolated *Ostertagia ostertagi* population. The CrIs were calculated using either *eggCounts* (EC) or *bayescount* (BC) models and interpreted based on either the revised guidelines (RG) for the faecal egg count reduction test (Kaplan et al., 2023) with corresponding 90 % CrIs, or on the original guidelines (OG) (Coles et al., 1992) with 95 % CrIs. Each point represents the mean FECR, with colour indicating the resistance status classified to the strongyle population: green, susceptible/normal; red, resistant; pink, low resistant / suspected resistant; orange, inconclusive / suspected susceptible.

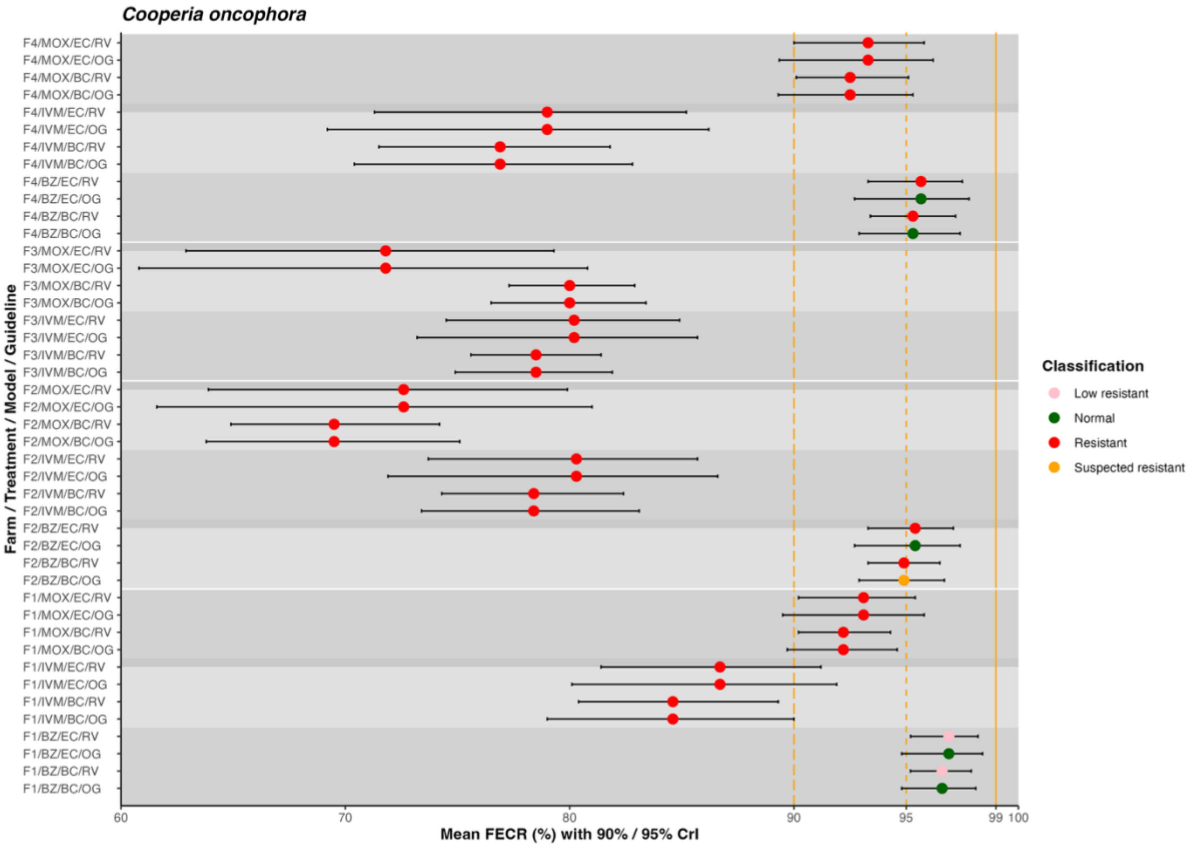


**Figure S2 Comparison of the *eggCounts* and *bayescount* faecal egg count reduction estimates for the interpolated *Cooperia oncophora* population**

Faecal egg count reductions (FECR) with the credible intervals (CrIs) for anthelmintic treatment against the interpolated *Cooperia oncophora* population. The CrIs were calculated using either *eggCounts* (EC) or *bayescount* (BC) models and interpreted based on either the revised guidelines (RG) for the faecal egg count reduction test (Kaplan et al., 2023) with corresponding 90 % CrIs, or on the original guidelines (OG) (Coles et al., 1992) with 95 % CrIs. Each point represents the mean FECR, with colour indicating the resistance status classified to the strongyle population: green, susceptible/normal; red, resistant; pink, low resistant / suspected resistant; orange, inconclusive / suspected susceptible.

### Supplementary File 5

**Table S3. The inter-rater agreement between *eggCounts* and *bayescount* results based on the revised guidelines for the faecal egg count reduction test (Kaplan et al., 2023) for the entire strongyle population.**

| ***bayescount*** | ***eggCounts*** | | | |
|---|---|---|---|---|
| | **Susceptible** | **Inconclusive** | **Low resistant** | **Resistant** |
| **Susceptible** | - | - | - | - |
| **Inconclusive** | - | - | - | - |
| **Low resistant** | 1 | - | 3 | - |
| **Resistant** | - | - | - | 7 |
| | Kohen's k = 0.656 (substantial agreement) | | | |

**Table S4. The inter-rater agreement between *eggCounts* and *bayescount* results based on the revised guidelines for the faecal egg count reduction test (Kaplan et al., 2023) for the interpolated *Ostertagia ostertagi* population.**

| ***bayescount*** | ***eggCounts*** | | | |
|---|---|---|---|---|
| | **Susceptible** | **Inconclusive** | **Low resistant** | **Resistant** |
| **Susceptible** | - | - | - | - |
| **Inconclusive** | - | - | - | - |
| **Low resistant** | 1 | - | 3 | - |
| **Resistant** | - | - | - | 7 |
| | Kohen's k = 0.656 (substantial agreement) | | | |

**Table S5. The inter-rater agreement between *eggCounts* and *bayescount* results based on the revised guidelines for the faecal egg count reduction test (Kaplan et al., 2023) for the interpolated *Cooperia oncophora* population.**

| ***bayescount*** | ***eggCounts*** | | | |
|---|---|---|---|---|
| | **Susceptible** | **Inconclusive** | **Low resistant** | **Resistant** |
| **Susceptible** | - | - | - | - |
| **Inconclusive** | - | - | - | - |
| **Low resistant** | - | - | 1 | - |
| **Resistant** | - | - | - | 10 |
| | Kohen's k = 1 (perfect agreement) | | | |

**Table S6. Inter-rater agreement between the original guidelines (Coles et al., 1992) and the revised guidelines (Kaplan et al., 2023) based on the faecal egg count reduction test for the entire strongyle population analysed using *eggCounts*.**

| **Original guidelines** | **Revised guidelines** | | | |
|---|---|---|---|---|
| | **Susceptible** | **Inconclusive** | **Low resistance** | **Resistant** |
| Normal | 1 | - | 3 | - |
| Suspected susceptibility | - | - | - | - |
| Suspected resistance | - | - | - | - |
| Resistance | - | - | - | 7 |
| | Cohen's k = 0.656 (substantial agreement) | | | |

**Table S7. Inter-rater agreement between the original guidelines (Coles et al., 1992) and the revised guidelines (Kaplan et al., 2023) based on the interpolated faecal egg count reduction test for *Ostertagia ostertagi* analysed using *eggCounts*.**

| **Original guidelines** | **Revised guidelines** | | | |
|---|---|---|---|---|
| | **Susceptible** | **Inconclusive** | **Low resistance** | **Resistant** |
| Normal | 1 | - | 3 | 2 |
| Suspected susceptibility | - | - | - | - |
| Suspected resistance | - | - | - | 1 |
| Resistance | - | - | - | 4 |
| | Cohen's k = 0.33 (fair agreement) | | | |

**Table S8. Inter-rater agreement between the original guidelines (Coles et al., 1992) and the revised guidelines (Kaplan et al., 2023) based on the interpolated faecal egg count reduction test for *Cooperia oncophora* analysed using *eggCounts*.**

| **Original guidelines** | **Revised guidelines** | | | |
|---|---|---|---|---|
| | **Susceptible** | **Inconclusive** | **Low resistance** | **Resistant** |
| Normal | - | - | 1 | 2 |
| Suspected susceptibility | - | - | - | - |
| Suspected resistance | - | - | - | - |
| Resistance | - | - | - | 8 |
| | Cohen's k = 0.19 (slight agreement) | | | |

### Supplementary file 5

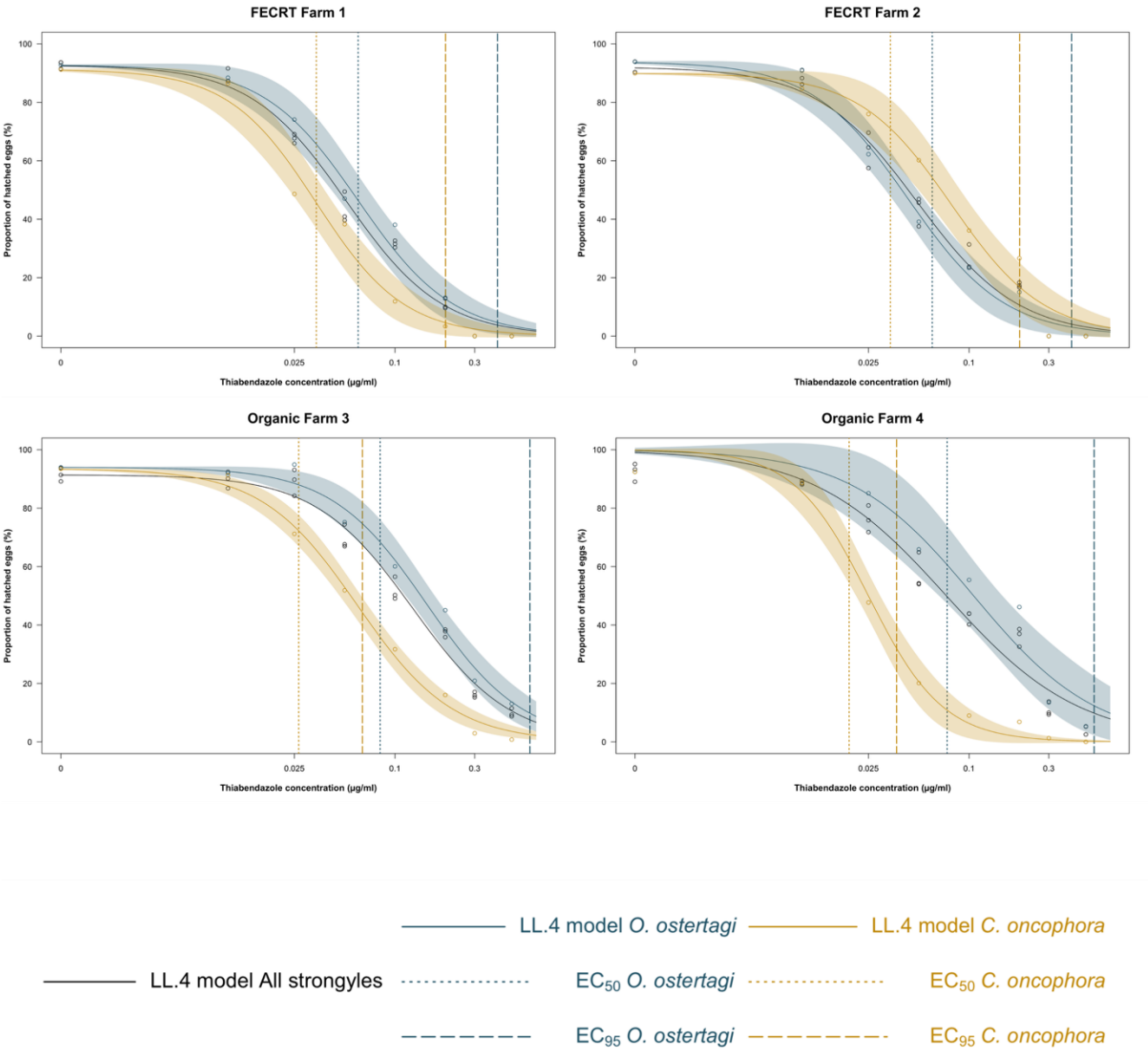


**Figure S3: Individual population dose-response curves**

The results of the egg hatch test performed with thiabendazole (TBZ) in 2023 on each study farm are presented at the extrapolated species and total strongyle level. Presented are the LL.4 model dose-response curves and the maximum effective concentrations (EC), $EC_{50}$ and $EC_{95}$ values.

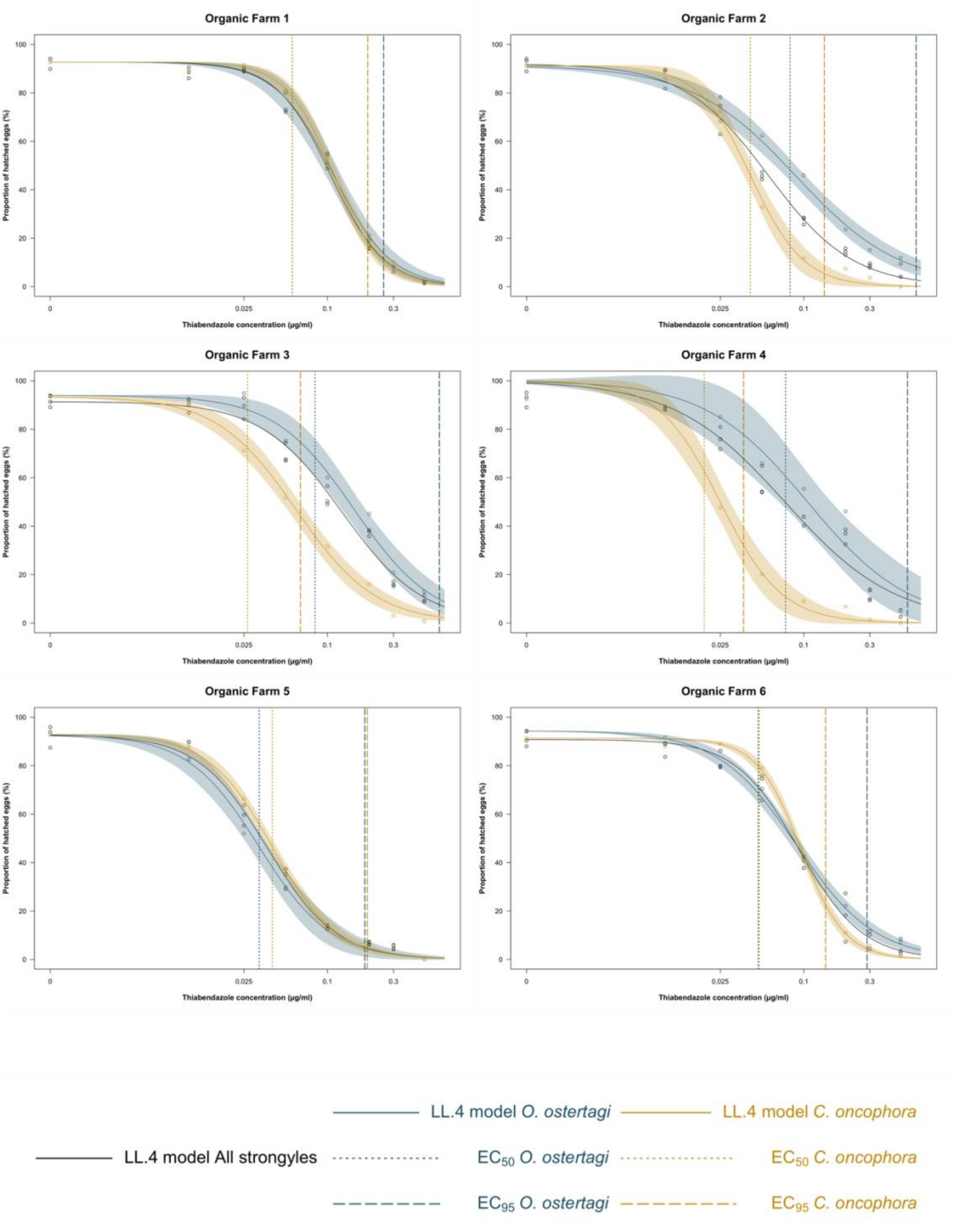


**Figure S3 (continued): Individual population dose-response curves**

The results of the egg hatch test performed with thiabendazole (TBZ) in 2023 on each study farm are presented at the extrapolated species and total strongyle level. Presented are the LL.4 model dose-response curves and the maximum effective concentrations (EC), $EC_{50}$ and $EC_{95}$ values.

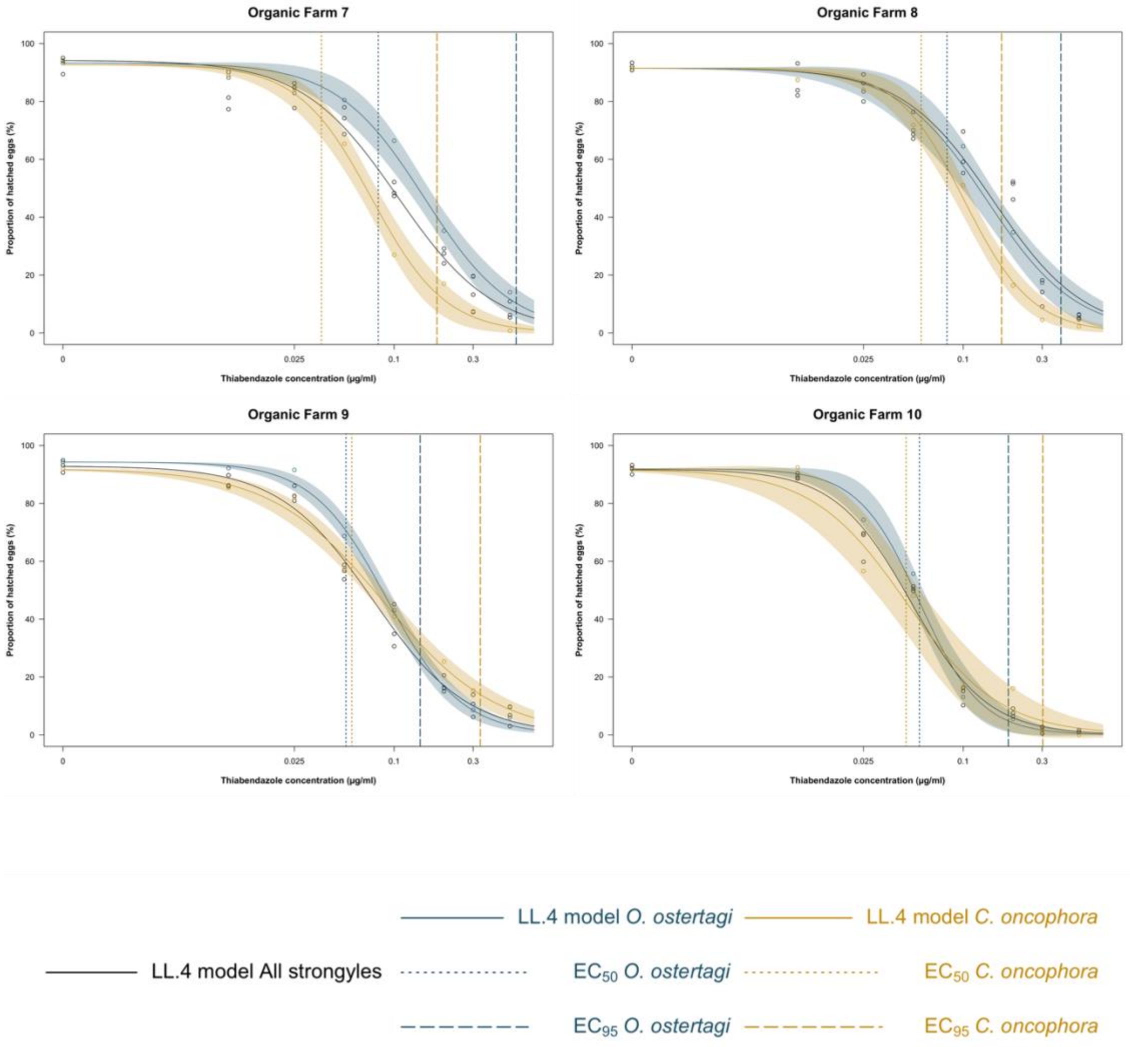


**Figure S3 (continued): Individual population dose-response curves**

The results of the egg hatch test performed with thiabendazole (TBZ) in 2023 on each study farm are presented at the extrapolated species and total strongyle level. Presented are the LL.4 model dose-response curves and the maximum effective concentrations (EC), $EC_{50}$ and $EC_{95}$ values.

### Supplementary File 6

**Table S8. Prevalence of gastrointestinal nematode species composition identified by mixed amplicon sequencing of the ITS-2 region in ten populations from seven Scottish dairy farms.**

| ID | *Os. ostertagi* | *C. oncophora* | *Tr. axei* | *Tr. colubriformis* | *Te. circumcincta* | *Os. leptospicularis* | *Oe. venulosum* | *Oe. radiatum* | *H. contortus* | *Tr. vitrinus* | *N. helvetianus* | *Trichostrongylus* spp. | *Cooperia* spp. |
|---|---|---|---|---|---|---|---|---|---|---|---|---|---|
| **FECRT_1/BZ/PRE** | 52.1 | 29.7 | 14 | 2.7 | 1.5 | 0 | 0 | 0 | 0 | 0 | 0 | 0 | 0 |
| **FECRT_1/BZ/POST** | 85.2 | 7.9 | 3 | 0 | 3.9 | 0 | 0 | 0 | 0 | 0 | 0 | 0 | 0 |
| **FECRT_2/IVM/PRE** | 67.2 | 21.3 | 6.8 | 2.7 | 0.3 | 0 | 0 | 0 | 1.1 | 0 | 0 | 0 | 0.6 |
| **FECRT_2/IVM/POST** | 51.7 | 39.9 | 0 | 0 | 2.7 | 0 | 0 | 0 | 0 | 0 | 0 | 0 | 5.7 |
| **FECRT_3/MOX/PRE** | 42.6 | 45.8 | 5.9 | 0 | 0 | 0 | 0 | 0 | 0 | 0 | 2.8 | 0 | 2.9 |
| **FECRT_3/MOX/POST** | 28 | 71.1 | 0 | 0 | 0 | 0 | 0 | 0 | 0 | 0 | 0 | 0 | 0.9 |
| **ORGANIC02** | 50.9 | 36 | 3.8 | 5.9 | 0 | 0 | 1.9 | 0.4 | 0 | 0 | 0.7 | 0.4 | 0 |
| **ORGANIC03** | 67 | 15 | 10.7 | 2.5 | 1.4 | 0 | 0 | 1.6 | 0.8 | 0.2 | 0 | 0.8 | 0 |
| **ORGANIC04** | 66.6 | 21 | 3.8 | 0.5 | 0 | 1.1 | 3.9 | 3.1 | 0 | 0 | 0 | 0 | 0 |
| **ORGANIC10** | 50.3 | 32.9 | 5 | 4.8 | 0 | 0 | 2.9 | 3.8 | 0 | 0 | 0 | 0.3 | 0 |
